# Dipeptidyl peptidase 9 triggers BRCA2 degradation by the N-degron pathway to promote DNA-damage repair

**DOI:** 10.1101/2020.08.24.265033

**Authors:** Maria Silva-Garcia, Oguz Bolgi, Breyan Ross, Esther Pilla, Vijayalakshmi Kari, Markus Killisch, Nadine Stark, Christof Lenz, Melanie Spitzner, Mark D. Gorrell, Marian Grade, Henning Urlaub, Matthias Dobbelstein, Robert Huber, Ruth Geiss-Friedlander

## Abstract

Dipeptidyl peptidase 9 (DPP9) is a serine protease cleaving N-terminal dipeptides preferentially post-proline with (patho)physiological roles in the immune system and cancer. Only few DPP9 substrates are known. Here we identify an association of human DPP9 with the tumour suppressor BRCA2, a key player in repair of DNA double-strand breaks that promotes the formation of RAD51 filaments. This interaction is triggered by DNA-damage and requires access to the DPP9 active-site. We present crystallographic structures documenting the N-terminal Met_1_-Pro_2_ of a BRCA2_1-40_ peptide captured in the DPP9 active-site. Mechanistically, DPP9 targets BRCA2 for degradation by the N-degron pathway, and promotes RAD51 foci formation. Both processes are phenocopied by BRCA2 N-terminal truncation mutants, indicating that DPP9 regulates both stability and the cellular stoichiometric interactome of BRCA2. Consistently, DPP9-deprived cells are hypersensitive to DNA-damage. Together, we identify DPP9 as a regulator of BRCA2, providing a possible explanation for DPP9 involvement in cancer development.

## Introduction

<u>D</u>i<u>p</u>eptidyl <u>p</u>eptidases <u>9</u> (DPP9) is an intracellular protease of the DPPIV family, that is critical for neonatal survival (Gall et al. 2013) with roles in the immune response (Geiss-Friedlander et al. 2009; Okondo et al. 2016; Okondo et al. 2018; Johnson et al. 2018; de Vasconcelos et al. 2019). Deregulation of DPP9 is connected to pathophysiological conditions such as tumorigenicity (Tang et al. 2017; Smebye et al. 2017; Johnson et al. 2018; Spagnuolo et al. 2013), whereby the underlying mechanisms are poorly understood.

On the molecular level, proteases of the DPPIV family are serine aminopeptidases that remove dipeptides from the N-terminus of substrates having a Pro or an Ala residue in the second position (NH_2_-XaaPro/Ala) (Waumans et al. 2015). DPP8 and DPP9 are the only known intracellular members of this family, both localize to the cytosol, the nucleus and associate with the plasma membrane (Ajami et al. 2004; Justa-Schuch et al. 2014; H. Zhang, Chen, et al. 2015). DPP9 is significantly more abundant than DPP8, and its depletion leads to a drastic reduction in the capacity of cells to process proline containing peptides (Geiss-Friedlander et al. 2009). In addition to peptides, several proteins were identified as DPP9 substrates (H. Zhang, Maqsudi, et al. 2015; Wilson et al. 2013). These include the tyrosine kinase Syk, a central kinase in the B cell receptor mediated signalling pathway. Previously we showed that processing of the Syk N-terminus by DPP9 leads to Syk degradation, linking DPP9 to the N-degron pathway (Justa-Schuch et al. 2016), a branch of the ubiquitin-proteasome pathway which targets proteins with destabilizing N-terminal residues for degradation (Varshavsky 2019). Although DPP9 binds directly to Syk, in cells this interaction is stabilized by <u>F</u>ilami<u>n A</u> (FLNA) (Justa-Schuch et al. 2016), a scaffold protein that localizes both to the cytoplasm and to the nucleus, where it interacts with multiple proteins thereby regulating signalling networks (Yue et al. 2013; Yue et al. 2009; Velkova et al. 2014; Yue et al. 2012).

Here, we asked whether additional FLNA partners would be substrates of DPP9. Thus, we inspected the N-terminus of known FLNA-binding proteins for a potential DPP9 cleavage sequence. This search highlighted the N-terminus of the human BRCA2 (Met-Pro-Ile), which was previously shown to bind to FLNA (Yuan & Shen 2001b). Alignment of multiple species shows that the potential DPP9 cleavage site in the N-terminus of BRCA2 is conserved in many mammals (Figure 1 - figure supplement 1). In some species, the proline is replaced by an alanine, or the isoleucine is replaced by a valine (Met-<u>Pro/Ala</u>-Ile/Val) (http://slim.icr.ac.uk/proviz/proviz.php?uniprot_acc=P51587). Also these sequences present potential DPP9-cleavage sites.

BRCA2 is critical for repair of DNA <u>d</u>ouble <u>s</u>trand <u>b</u>reaks (DSBs) by the homologous recombination (HR) pathway (Moynahan et al. 2001). Such lesions present the most harmful form of DNA damage, as they can lead to genome rearrangement and genome instability if not correctly repaired. In contrast to the repair of DSBs by the non-homologous end-joining pathway which is error prone, the repair of such breaks by HR is mostly error free. In short, the repair of DSBs by the HR pathway starts with the generation of a 3’ single strand DNA (ssDNA) overhang by a process called 5’ end-resection. The ssDNA is subsequently covered by protein filaments of the strand-exchange protein RAD51. These filaments form a so-called displacement loop (D-loop), which invades into homologous sequences preferentially in the sister chromatid that serves as template for repair of the lesions by DNA polymerases (Scully et al. 2019; C.-C. Chen et al. 2018). The formation of the RAD51 filament is directly regulated by BRCA2, which interacts with multiple copies of RAD51 via eight so called BRC repeats located in the centre of BRCA2 and an additional RAD51 binding domain at the C-terminus (A. A. Davies et al. 2001; Pellegrini et al. 2002; Esashi et al. 2007; O. R. Davies & Pellegrini 2007). By simultaneously binding multiple monomers of RAD51, BRCA2 provides a rapid mechanism for loading and assembly of RAD51 as filaments on the ssDNA (Thorslund et al. 2010; Jie Liu et al. 2010; Jensen et al. 2010; Carreira & Kowalczykowski 2011; Shahid et al. 2014; Sánchez et al. 2017). The importance of BRCA2 is highlighted by the fact that it is often mutated in patients with familial breast and ovarian cancers (Scully et al. 2019; C.-C. Chen et al. 2018). Intriguingly, although critical for DSB repair, BRCA2 steady-state protein levels were shown to be lower following DNA damage, a process that is counteracted by the ubiquitin-specific protease 21 (USP21) and by proteasome inhibition (Schoenfeld et al. 2004; Jinping Liu et al. 2017).

Given the interactions of DPP9 and BRCA2 with FLNA (Justa-Schuch et al. 2016; Yuan & Shen 2001a) and the presence of a conserved putative DPP9 cleavage site in an unstructured N-terminal sequence of BRCA2, we asked whether BRCA2 is regulated by DPP9. In this work we show that DPP9 promotes BRCA2 degradation and also activity. We suggest that DPP9 is critical for establishing the cellular stoichiometric interactome of BRCA2.

## Results

### DNA damaging conditions trigger an interaction between BRCA2 and DPP9, requiring DPP9 active site

As a starting point we asked whether DPP9 and BRCA2 interact in cells. Thus, we performed Proximity Ligation Assays (PLA), a highly sensitive method that is especially suited for detection and quantification of dynamic interactions within cells. Initial analysis of HeLa cells did not detect specific PLA signals for BRCA2-DPP9 above the background signals that are observed in control cells silenced for either DPP9 or BRCA2 (Figure 1 – figure supplement 2A and 2B). However, since BRCA2 is critical for repair of DSBs, these assays were also carried out in cells that were exposed to Mitomycin C (MMC), a chemotherapeutic agent that forms inter-strand DNA crosslinks which are converted to secondary DSBs. Strikingly, whereas MMC did not trigger a close proximity between DPP9 and BRCA2 in cells silenced for FLNA, specific signals for DPP9-BRCA2 PLA were detected both in the cytoplasm and in the nucleus of mock-silenced cells in response to MMC (Figure 1A and 1B, Figure 1 – figure supplement 2A).

**Figure 1.**
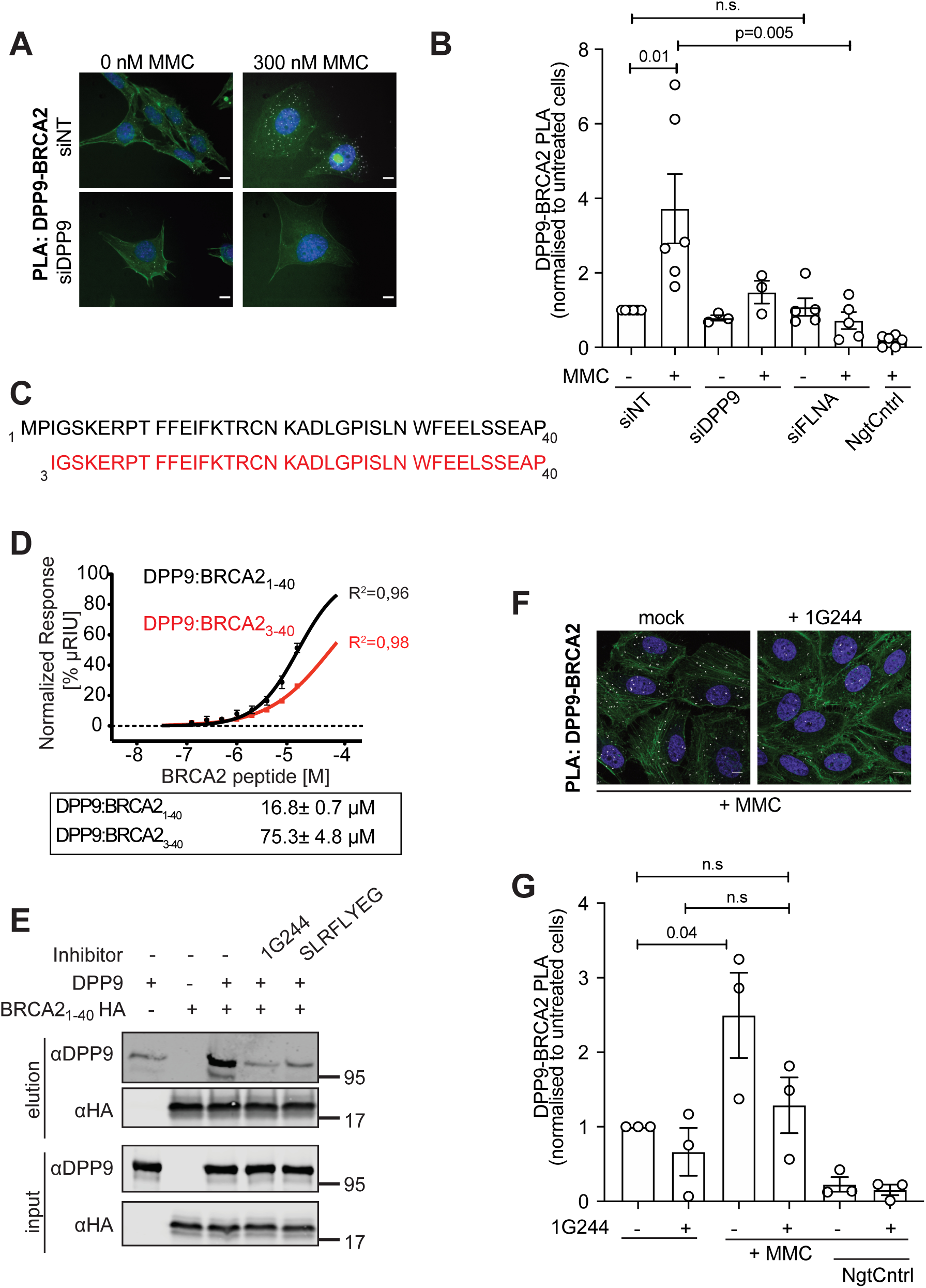
DNA damage triggers an interaction between DPP9 and BRCA2, requiring access to DPP9 active site. (A) Proximity ligation assay (PLA) images showing that MMC triggers an interaction between endogenous DPP9 and BRCA2 in HeLa WT cells. Each white dot represents a single PLA event between DPP9 and BRCA2. Phalloidin (green) stains actin filaments, DAPI (blue) stains the nucleus. Scale bar 10 µm. (B) Graph summarizing independent biological replicates of the PLA for BRCA2-DPP9. In each replicate, more than 125 cells were quantified for each condition respectively. PLA of the technical control samples (NgtCntrl) did not include the BRCA2 antibody. Each point represents the mean of one replicate, normalized to signals in control HeLa cells (siNT -MMC). Mean ± SEM, data were analysed by a two-way ANOVA, with Tukey’s Multiple Comparison test. (C) Amino acid sequences of synthetic peptides corresponding to the N-terminus of intact BRCA2 (BRCA2_1-40_) and truncated (BRCA2_3-40_). (D) Surface Plasmon Resonance (SPR) data showing a direct interaction of purified DPP9 with the BRCA2_1-40_ peptide. Binding affinity was significantly reduced for the truncated BRCA2_3-40_ peptide, which lacks the DPP9 cleavage site (Met-Pro). A serial dilution of BRCA2-derived peptides was injected over a surface covered with DPP9. Equilibrium binding isotherms obtained for interactions measured between DPP9 and BRCA2_1-40_ (black line) and BRCA2_3-40_ (red line). Data were fitted to a sigmoidal dose-response curve fit. Shown are mean ± SEM of the triplicate. Box summarizes the calculated affinities (KD values). (E) Direct binding of an N-terminal BRCA2 fragment to the relaxed form of DPP9. Recombinant fragments corresponding to the N-terminus of intact BRCA2 with a C-terminal HA tag (BRCA2_1-40_HA) were immobilized on HA-beads and tested for binding to purified DPP9. Binding of DPP9 to the BRCA2 N-terminal fragment was impaired in the presence of competitive DPP9 inhibitors (1G244 or SLRFLYEG). Representative data of three independent experiments is shown. (F) Fewer DPP9-BRCA2 PLA interactions (white dots) in HeLa cells treated with 1G244, which blocks the access to the DPP9 active site. Control cells were mock treated with DMSO. Scale bar 10 µm. Phalloidin (green) stains actin filaments, DAPI (blue) stains the nucleus. (G) Graph summarizing 3 independent biological replicates of the PLA for BRCA2-DPP9. In each replicate, more than 50 cells were quantified for each condition respectively. PLA of the technical control samples (NgtCntrl) did not include the BRCA2 antibody. Each point represents the mean of a single replicate, normalized to signals in control HeLa cells (siNT - 1G244). Shown are Mean ± SEM, data were analysed by a two-way ANOVA, with Tukey’s Multiple Comparison test.

**Figure 1 - figure supplement 1:**
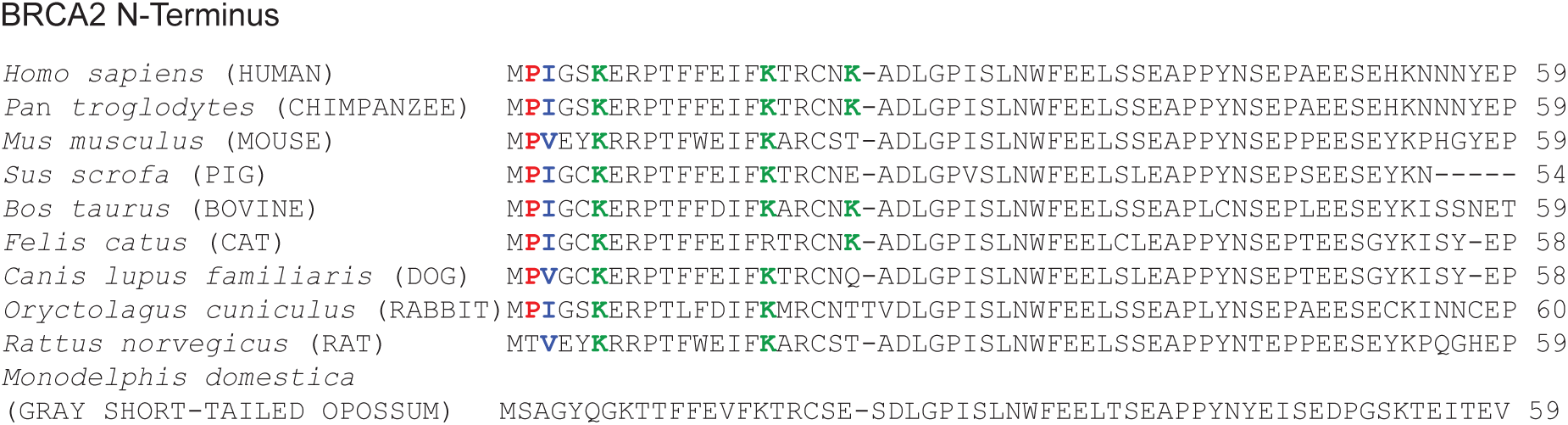
A potential DPP9 cleavage site in BRCA2. Alignment of the BRCA2 N-terminus in several species. The potential DPP9 cleavage site (Met-Pro↓Ile/Val) is shown in bold. The alignment was performed with Clustal Omega (multiple sequence alignment) program. Conserved lysine residues are highlighted in green.

**Figure 1 - figure supplement 2:**
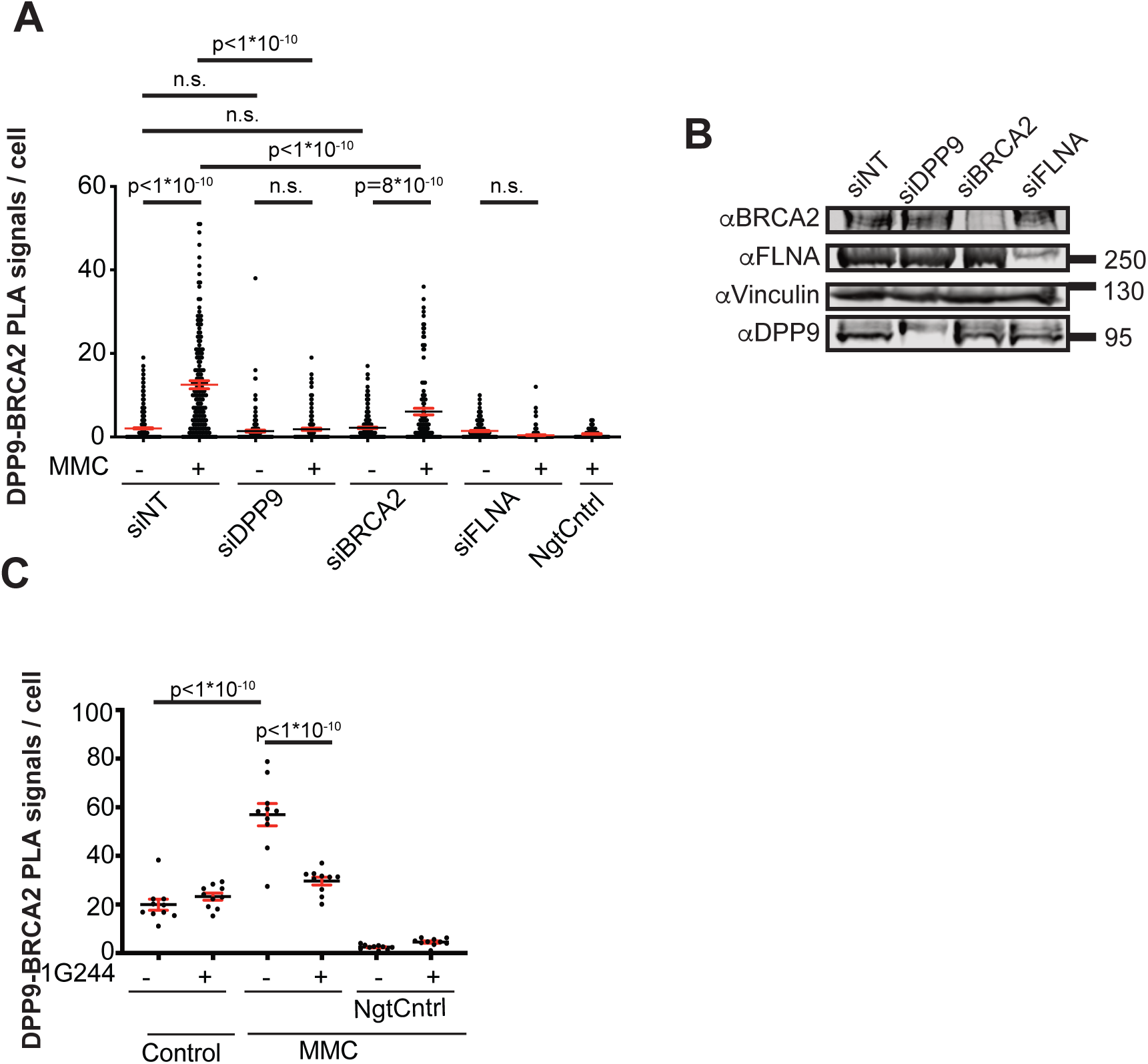
DNA damage triggers an interaction between DPP9 and BRCA2, requiring access to DPP9 active site. (A) Quantification of one PLA experiment a showing close proximity between DPP9 and BRCA2 that is triggered by MMC. PLA signals from more than 125 cells were quantified in each condition respectively. Mean ± SEM, data were analysed by a two-way ANOVA, with Tukey’s Multiple Comparison test. (B) Western Blots of the corresponding silenced cells (without MMC) analysed by PLA (A). For control, cells were silenced for DPP9 (siDPP9), BRCA2 (siBRCA2), FLNA (siFLNA) or treated with non-targeting siRNA (siNT). (C) Quantification of one of three PLAs, showing that the interaction between DPP9 and BRCA2 is blocked in the presence of the inhibitor 1G244. In each experiment, signals from more than 50 cells per condition per assay were quantified. Mean ± SEM, data were analysed by a two-way ANOVA, with Tukey’s Multiple Comparison test.

Subsequently, to examine whether the interaction between DPP9 and BRCA2 involves the N-terminus of BRCA2, Surface Plasmon Resonance (SPR) assays were performed. This approach documented direct binding of purified recombinant DPP9 to a peptide corresponding to the first forty amino acids of BRCA2 (BRCA2_1-40_), with a binding affinity (KD) of 16.8±0.7 µM (Figure 1C and 1D). A truncated form of the synthetic BRCA2 fragment, lacking the N-terminal residues Met_1_-Pro_2_ (BRCA2_3-40_), bound to DPP9 with lower affinity (KD) of 75.3± 4.8 µM, highlighting the importance of the N-terminal dipeptide Met_1_-Pro_2_ for binding to DPP9 (Figure 1D).

Consistent with the SPR measurements, pull-down assays confirmed a direct binding between DPP9 and a construct corresponding to the first forty amino acids of BRCA2 fused to a C-terminal HA tag (BRCA2_1-40_HA) (Figure 1E). These assays were also performed in the presence of SLRFLYEG or 1G244, two competitive inhibitors of DPP9 (Wu et al. 2009; Pilla et al. 2013). Previously, we had shown that binding of 1G244 or SLRFLYEG, initiates a rearrangement and a closed conformation of the DPP9 active site (Ross et al. 2018). Importantly, the binding of DPP9 to BRCA2_1-40_HA was clearly blocked by SLRFLYEG and 1G244, demonstrating the importance of the open DPP9 active site confirmation for binding to the BRCA2 N-terminus (Figure 1E). The significance of access to the active site of DPP9 was further challenged in cells. Consistent with the results obtained with the recombinant factors, cells treated with 1G244 did not show a significant increase in DPP9-BRCA2 PLA events in response to MMC (Figure 1F and 1G, Figure 1 – figure supplement 2C). Taken together, these data show an interaction between DPP9 and BRCA2 that is triggered by DNA damage and requires access to the DPP9 active site, a characteristic for interactions between an enzyme and its substrate.

### DPP9 removes the BRCA2 N-terminal dipeptide Met_1_Pro_2_

Since the interaction with BRCA2 requires access to the active site of DPP9, we next tested whether DPP9 can hydrolyse BRCA2 N-terminal peptide. *In vitro* reactions were carried out with purified DPP9 and the BRCA2_1-40_ N-terminal peptide followed by mass spectrometry analysis. Control reactions contained no DPP9, or included the DPP9 inhibitor SLRFLYEG. These assays demonstrated processing of BRCA2_1-40_ N-terminal peptide by purified DPP9, leading to removal of the dipeptide Met_1_Pro_2_, with the resulting product (BRCA2_3-40_) having an isoleucine in the neo N-terminus (Figure 2A, Met_1_Pro_2_↓Ile_3_).

**Figure 2.**
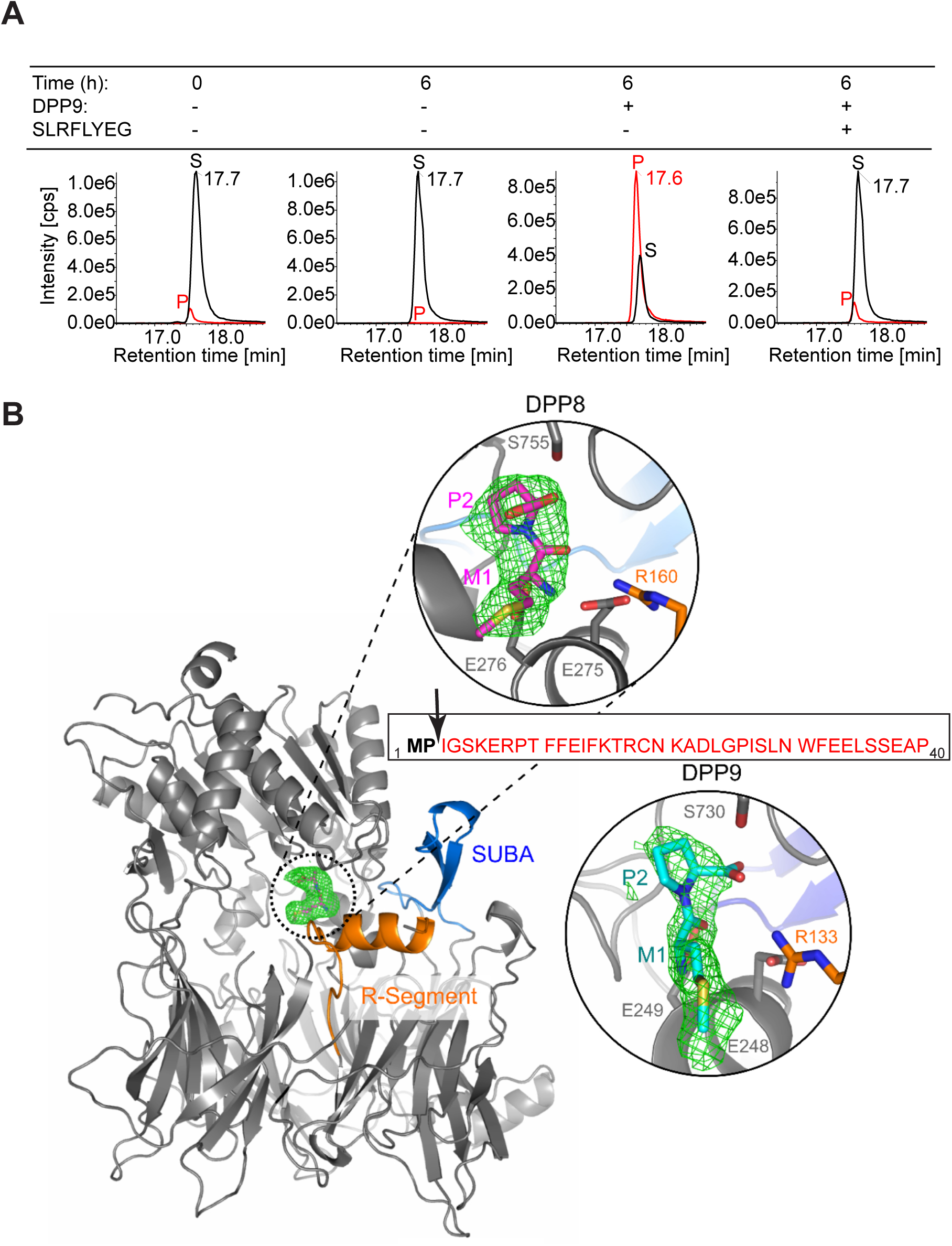
DPP9 enzymatic activity cleaves the BRCA2_1-40_ N-terminal peptide. (A) A representative experiment showing in vitro processing of BRCA2_1-40_ synthetic peptide by purified recombinant DPP9. BRCA2_1-40_ peptide was incubated with DPP9 for 6 hours. Control samples included the DPP9 inhibitor SLRFLYEG. Samples were analysed by high-resolution liquid chromatography/tandem mass spectrometry, in quadruplicate. The panels show extracted MS1 ion chromatograms for both substrate BRCA2_1-40_ peptide (MPIGSKERPT…) (labelled S, [M+5H]5+ m/z 917.8637; retention time 17.7 minutes) and product BRCA2_3-40_ peptide (IGSKERPT…) (labelled P, [M+5H]5+ m/z 872.2451; retention time 17.6 minutes). The identity of the product peak was established both by accurate mass measurement to within 5 ppm and by product ion spectra (data not shown). (B) Shown is the N-terminal di-peptide of BRCA2 bound to both DPP9 and to DPP8. Omit map (Fo-Fc; 3σ) of a BRCA2 peptide soaked in DPP8 crystals (C222_1_) and in DPP9 crystals (P12_1_1). The R-Segment and SUBA are highlighted in orange and blue, respectively. The arrow marks the position where the BRCA2 peptide is cleaved. For simplification, shown here is a monomer of DPP8. Both zoomed views are rotated 45° with respect to the monomer view to better display the ligand.

To further understand the cleavage event, DPP9 crystals were soaked with the BRCA2_1-40_ peptide. Strikingly, DPP9 crystals exhibited positive electron density at the active site corresponding to the N-terminal Met_1_Pro_2_ di-peptide with the proline residue occupying the S1 subsite. Since DPP8 and DPP9 share a very similar substrate binding mechanisms *in vitro*, and since DPP9 crystals are notorious for being less well ordered (Ross et al. 2018), we also tested for a possible interaction between DPP8 and BRCA2. Similarly, in DPP8 all 3 copies of the asymmetric unit exhibit clearly interpretable electron density for the di-peptide. The well-ordered R-segment, a hallmark of ligand binding, was well defined and ordered and supports full occupation of the active sites. There was no trace of electron density for the rest of the peptide (Figure 2B and Table 1). These structures with the di-peptide captured in the active site clearly document intra-crystalline enzymatic activity of DPP8 and DPP9 resulting in the hydrolysis of the BRCA2_1-40_ N-terminal peptide, and release of the BRCA2_3-40_ peptide. Together with the cellular assays, we conclude that DPP9 processes the N-terminus of BRCA2.

**Table 1:** Crystallographic parameters for DPP8 and DPP9 -BRCA2 peptide bound structures.

|  | DPP8 - BRCA2 peptide | DPP9 - BRCA2 peptide |
| --- | --- | --- |
| **Data collection** |  |  |
| Space group | C222<sub>1</sub> | P12<sub>1</sub>1 |
| Resolution (Å) | 44.58 - 3.20 | 43.47 - 3.00 |
|  | (3.28 - 3.20)<sup>a</sup> | (3.19 - 3.00)<sup>a</sup> |
| a, b, c (Å) | 162.73 246.07 261.81 | 119.52 117.22 163.40 |
| $\alpha$, $\beta$, $\gamma$ (°) | 90, 90, 90 | 90, 105.74, 90 |
| R<sub>meas</sub> (%) | 14.3 (139.9)<sup>a</sup> | 15.9 (37.6)<sup>a</sup> |
| CC<sub>1/2</sub> (%) | 99.8 (73.6)<sup>a</sup> | 98.9 (93.9)<sup>a</sup> |
| I/σ(I) | 17.2 (1.8)<sup>a</sup> | 5.9 (3.0)<sup>a</sup> |
| Completeness (%) | 99.9 (99.2)<sup>a</sup> | 87.8 (88.8)<sup>a</sup> |
| Multiplicity | 8.37 (8.34)<sup>a</sup> | 2.65 (2.53)<sup>a</sup> |
| Total observations | 725357 (52815)<sup>a</sup> | 202589 (33069)<sup>a</sup> |
| Total unique observations | 86450 (6331)<sup>a</sup> | 76534 (13092)<sup>a</sup> |
| **Refinement** |  |  |
| R<sub>cryst</sub>/R<sub>free</sub> (%) | 18.39/22.47 | 28.5/34.5<sup>b</sup> |
| Number of reflections | 82126 (4323)<sup>c</sup> | 72706(3827)<sup>c</sup> |
| R.m.s.d. bond lengths (Å) | 0.003 | 0.004 |
| R.m.s.d. bond angles (°) | 0.984 | 1.120 |
| Number of atoms | 20409 | 26316 |
| Average B-factor (Å<sup>2</sup>) | 84.8 | 25.5 |
| **Ramachandran plot (%)** |  |  |
| Preferred region | 93.4 | 91.9 |
| Allowed region | 5.9 | 7.2 |
| Outliers | 0.6 | 0.8 |
| <sup>a</sup> Values in parentheses correspond to the highest-resolution shell |  |  |
| <sup>b</sup> The relatively high values are due to the crystals displaying multiple splits (Ross et al. 2018) |  |  |
| <sup>c</sup> Values in parentheses correspond to free R-value test set (5%) |  |  |

### DPP9-deficient cells are hypersensitive to genotoxic agents

The above data show that MMC-induced DNA damage triggers an interaction between DPP9 and BRCA2, which requires access to the active site of DPP9, and more importantly, that DPP9 cleaves BRCA2 N-terminus. Since BRCA2 is critical for repair of DSBs by HR, we raised the question whether DPP9 is relevant for repair of DNA damage. A common feature for cells with defects in repair of DSBs, is a hypersensitivity to DNA-damaging agents. Thus, the response of DPP9-deficient cells to the presence of genotoxic agents was assessed by analyzing the viability of these cells and their capacity to form colonies.

To test for the importance of DPP9, we assayed HeLa DPP9 KD cells that express lower DPP9 levels (50%) due to constitutive silencing of DPP9 (Figure 3 - figure supplement 1A and 1B, cells described first in (Justa-Schuch et al. 2016)). Strikingly, cell viability assays revealed that HeLa DPP9 KD cells are significantly more sensitive to MMC compared to the corresponding HeLa WT cells (Figure 3A). Subsequently, we tested the viability of the HeLa DPP9 KD cells following exposure to Olaparib, an FDA approved-drug that is used for treatment of germline BRCA1 and BRCA2-mutated metastatic breast cancers. Olaparib is an inhibitor of poly(ADP-ribose) polymerase (PARP), which plays a role in base excision repair of single-strand DNA (ssDNA) breaks. Cells which are defective in HR are hypersensitive to Olaparib because the accumulating ssDNA breaks are converted in cells to DSBs. Notably, we find that HeLa DPP9 KD cells are significantly more sensitive to Olaparib compared to control cells (Figure 3B), thus displaying a similar pattern to previous studies showing the sensitivity of BRCA2-deficient cells to Olaparib treatment (Menear et al. 2008; Rottenberg et al. 2008; Lord & Ashworth 2016).

**Figure 3.**
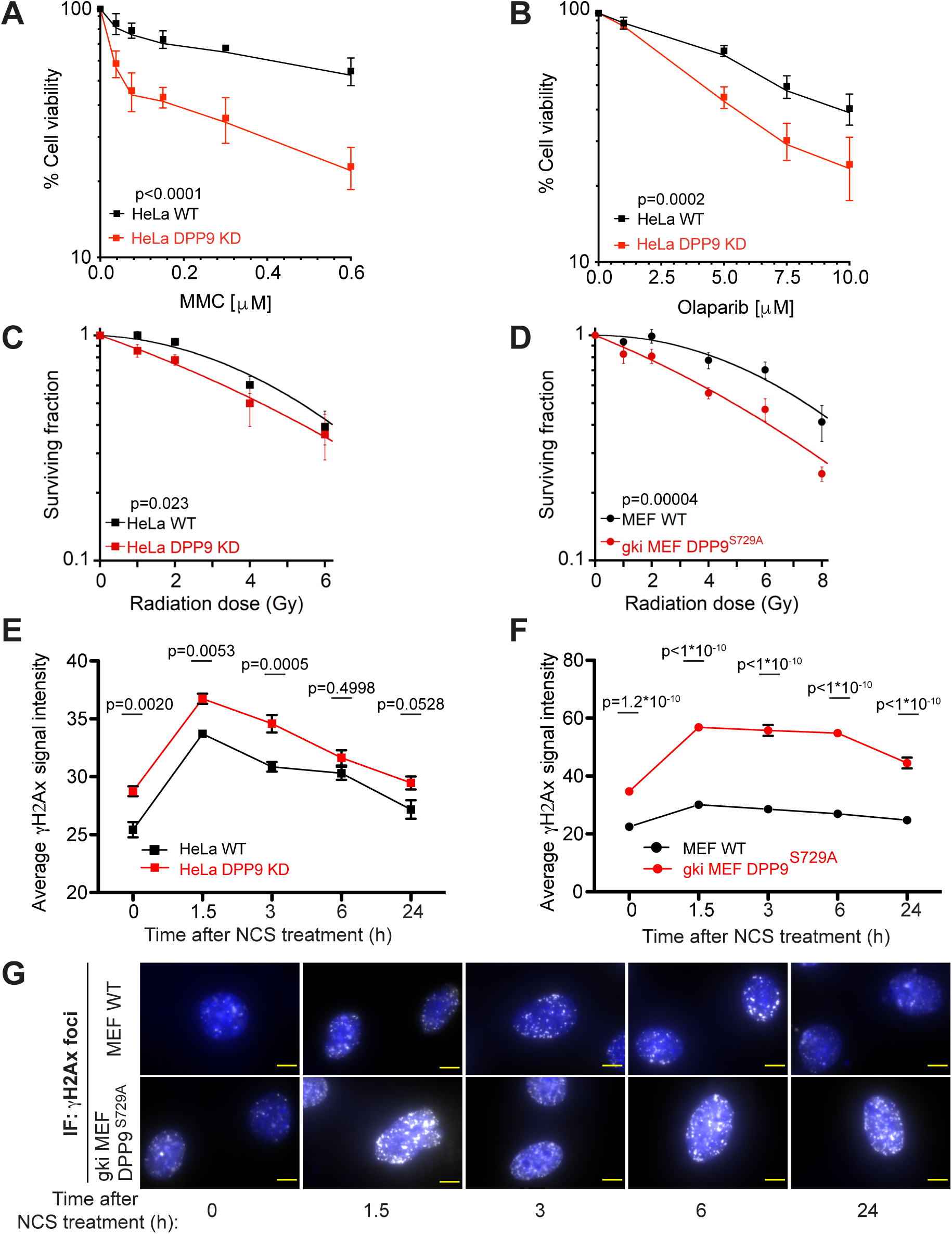
Cells with reduced DPP9 activity are hypersensitive to genotoxic agents. (A and B) Dose-dependent viability assays showing a higher sensitivity of HeLa DPP9 KD cells to (A) MMC and (B) Olaparib. Mean ± SEM of three independent biological replicates, with each experiment performed in triplicate. Data were analysed by an unpaired two-way ANOVA with Sidak’s multiple comparison test. (C and D) Quantification of colonies formed after γ-radiation of (C) HeLa WT and DPP9 KD, as well as (D) MEF WT and gki MEF DPP9^S729A^ cells, showing the mean ± SEM of the survival fraction (SF) from three independent experiments. Data were analysed by an unpaired two-way ANOVA. (E and F) Graphs showing that (E) HeLa DPP9 KD and (F) gki MEF DPP9^S729A^ cells accumulated more γH2AX signals in response to Neocarzinostatin (NCS) treatment compared to the corresponding control cells. All cells were treated with 250 ng/mL NCS for 30 minutes and allowed to recover for the indicated time-points. Data were analysed by an unpaired two-way ANOVA with Sidak’s multiple comparison test. (E) Signals from more than 1300 cells were quantified per condition per experiment. Mean ± SEM from four independent experiments. (F) Signals from more than 1700 cells were quantified per condition per experiment. Mean ± SEM from six independent experiments. (G) Representative immunofluorescence images showing the accumulation and recovery of γH2AX signals (white) in control MEF cells and in gki MEF DPP9^S729A^ cells following NCS treatment. The nucleus is shown in blue (DAPI). Scale bar 10 µm.

**Figure 3 - figure supplement 1:**
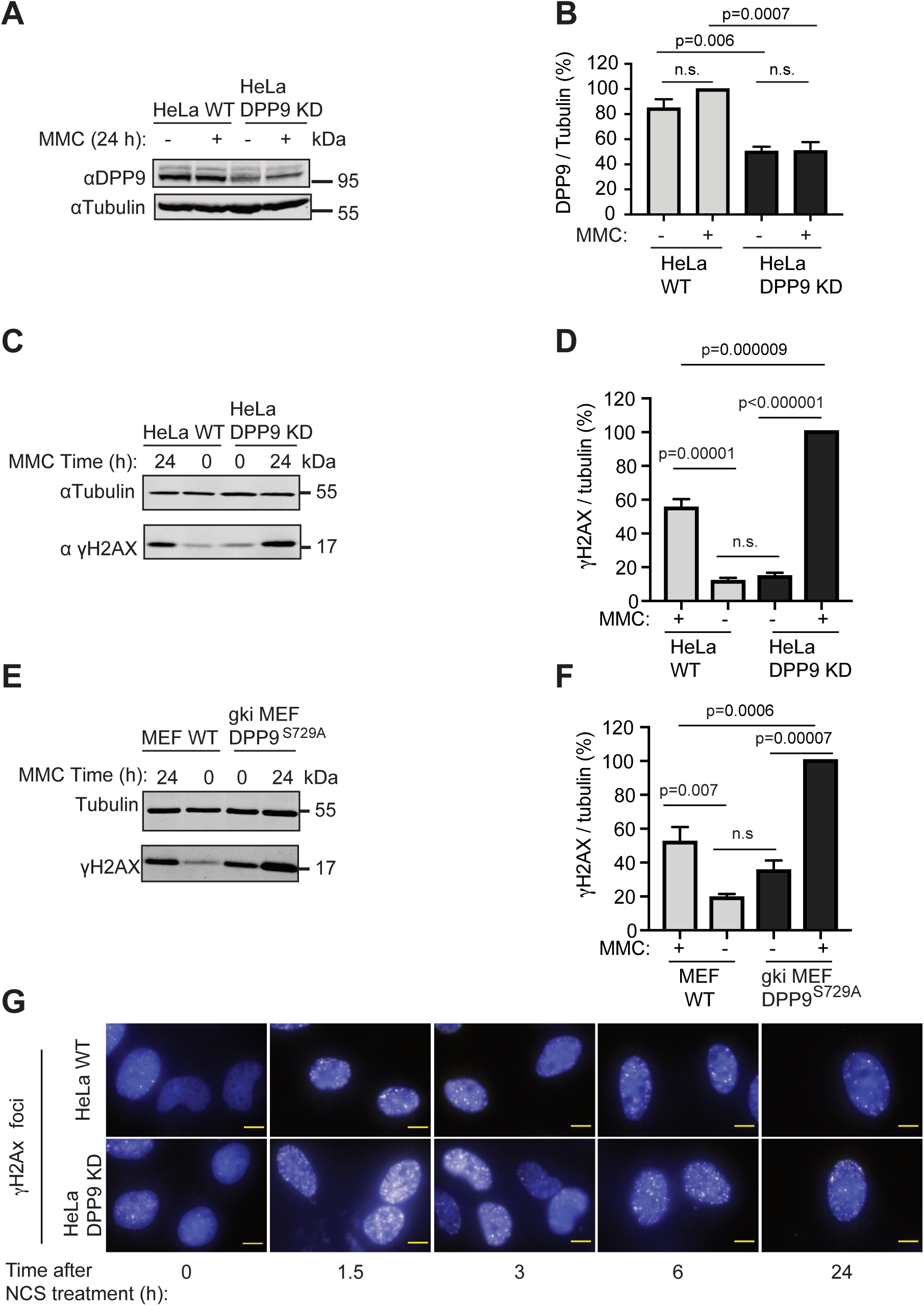
high γH2AX levels in DPP9-deprived cells. (A) Western-blot showing that the protein levels of DPP9 in HeLa DPP9 KD cells are significantly lower than those in the corresponding HeLa WT cells. DPP9 protein levels are not altered following exposure to 300 nM MMC for 24 hours. Tubulin was analysed as loading control. Representative images of one from three independent experiments are shown. (B) Quantification of DPP9 signals in HeLa WT and DPP9 KD cells following exposure to 300 nM MMC for 24 hours. Tubulin was analysed as loading control. Mean ± SEM. Data were analysed by a two-way ANOVA, with Tukey’s Multiple Comparison test. (C) Higher γH2AX signals in HeLa DPP9 KD cells following exposure to 300 nM MMC for 24 hours. Tubulin was analysed as loading control. (D) Quantification of the γH2AX signals in HeLa DPP9 KD cells and HeLa WT cells described in (C). Mean ± SEM. Data were analysed by a two-way ANOVA, with Tukey’s Multiple Comparison test. (E) Higher γH2AX signals in gki MEF DPP9^S729A^ cells expressing enzymatically inactive DPP9 compared to MEF WT control cells following exposure to 300 nM MMC for 24 hours. Tubulin was analysed as loading control. (F) Quantification of the γH2AX signals in gki MEF DPP9^S729A^ cells and MEF WT cells described in (E). Mean ± SEM. Data were analysed by a two-way ANOVA, with Tukey’s Multiple Comparison test. (G) Accumulation and reduction of γH2AX signals (white) in the nuclei of HeLa DPP9 KD and HeLa WT cells following treatment with 250 ng/mL NCS for 30 minutes. Cells were allowed to recover for the specified time-points and then fixed. Shown are representative images of one from three independent experiments. γH2AX is shown in white, the nucleus is shown in blue. Scale bar 10 µm.

Furthermore, we examined the capacity of the HeLa DPP9 KD cells to form colonies following exposure to ionizing radiation (γ-radiation) as a source for DSBs. Colony formation assays revealed that HeLa DPP9 KD cells are significantly more sensitive to γ-radiation in comparison to the corresponding control cells (Figure 3C). To test for the importance of DPP9 activity we analysed the DPP9 gene knock-in mouse embryonic fibroblasts (MEFs), which express an enzymatically inactive DPP9 variant carrying a point mutation in its catalytic serine (gki MEF DPP9^S729A^ cells were first described in (Gall et al. 2013)). Similar to the HeLa DPP9 KD cells, also the MEF DPP9^S729A^ were significantly more sensitive to γ-radiation compared to the corresponding MEF WT cells (Figure 3D).

To visualize the presence of DSBs, we followed the phosphorylation of H2AX on serine 139 (γH2AX). This phosphorylation step is an early and decisive step in repair of DSBs, that marks the surrounding of the break and is used as a platform that initiates the recruitment of repair proteins (Turinetto & Giachino 2015). In line with the observed reduced viability, the HeLa DPP9 KD cells also accumulated significantly higher levels of γH2AX in response to MMC (Figure 3 - figure supplement 1C and 1D). Consistently, significantly higher γH2AX were observed in the gki MEF DPP9^S729A^ compared to the MEF WT in response to MMC (Figure 3 - figure supplement 1E and 1F). We further examined the capability of DPP9-deprived cells to recover from the DNA damage, by following the disappearance of γH2AX. As a source for DNA damage, cells were treated with the radiomimetic antibiotic Neocarzinostatin (NCS). Following a 30 minute incubation, NCS was removed and the levels of γH2AX foci were monitored over time. These assays revealed higher γH2AX levels in HeLa DPP9 KD cells compared to the corresponding control cells (Figure 3E, Figure 3 - figure supplement 1G). Importantly, significantly higher γH2AX levels were also present in gki MEF DPP9^S729A^ (Figure 3F and 3G).

### DPP9 targets BRCA2 for degradation following DNA damage

The results above show that cells-deprived of DPP9 are hypersensitive to genotoxic agents and accumulate more DSBs, indicating that the activity of DPP9 is critical for the capacity of cells to survive in response to DSBs. Since DPP9 interacts and cleaves BRCA2 N-terminus, we inspected more closely the effect of DPP9 on BRCA2. As a first step, we analysed for the interaction of BRCA2 with the <u>P</u>artner <u>a</u>nd localizer of <u>B</u>RCA<u>2</u> (PALB2), a protein that ensures the correct intra-nuclear localization of BRCA2 following DNA-damage (Xia et al. 2006; F. Zhang et al. 2009; Oliver et al. 2009).

In line with published data, we detected PLA signals documenting interactions between endogenous BRCA2 and PALB2 (Figure 4A and 4B, Figure 4 - figure supplement 1). Strikingly, significantly more BRCA2-PALB2 PLA events were present in cells silenced for DPP9, in the presence of MMC (Figure 4A and 4B, Figure 4 - figure supplement 1), suggesting that DPP9 limits the BRCA2-PALB2 interaction. Furthermore, we observed slightly higher steady state levels of BRCA2 in cells silenced for DPP9, raising the question whether DPP9 influences BRCA2 stability (Figure 4 - figure supplement 1).

**Figure 4.**
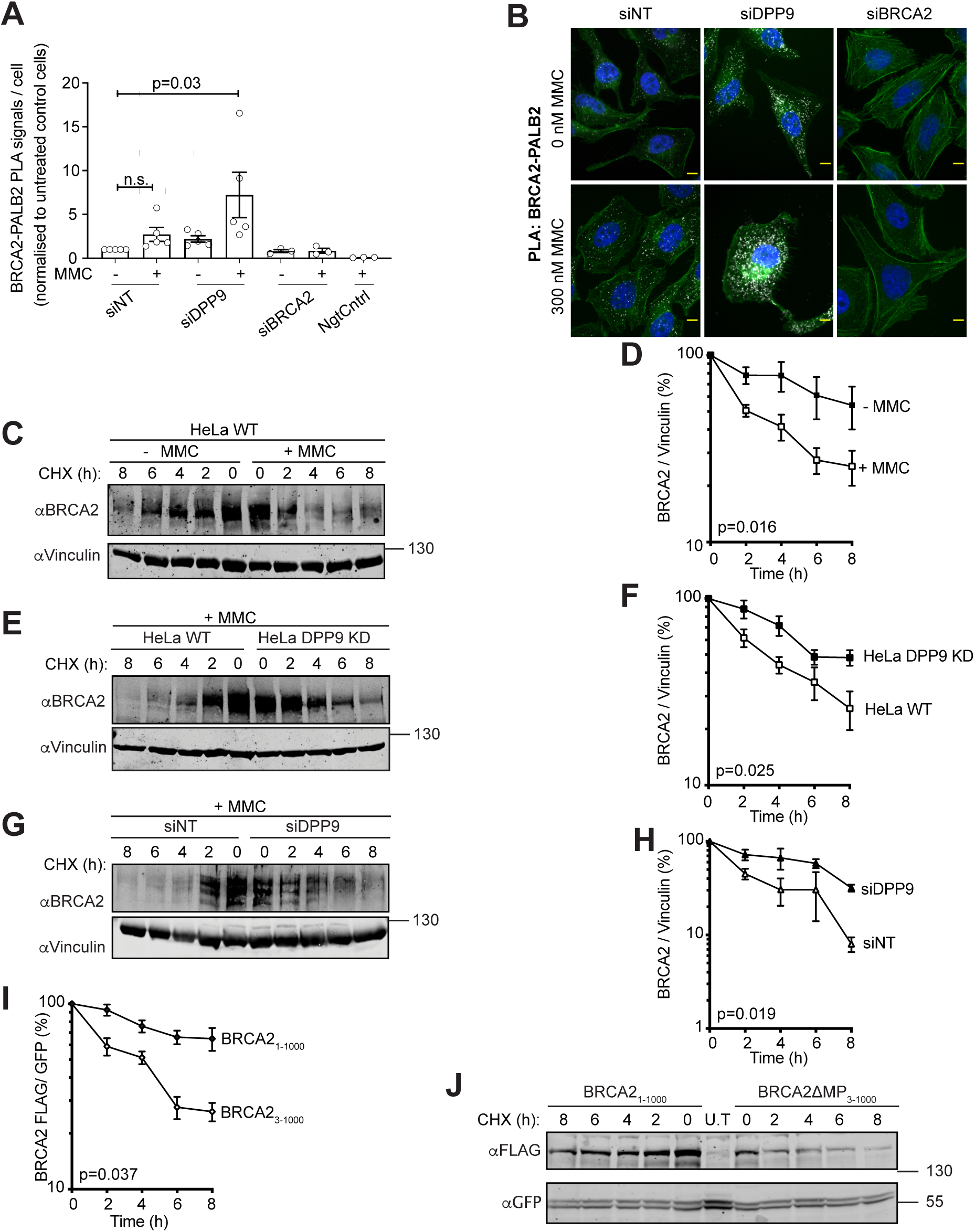
DPP9 triggers BRCA2 degradation in response to MMC. (A) PLA quantification showing more BRCA2-PALB2 events in HeLa cells silenced for DPP9 compared to cells treated with non-targeting siRNA (siNT). The graph summarizes five independent biological replicates. In each replicate, more than 100 cells were quantified for each condition respectively. PLA of the technical control samples (NgtCntrl) did not include the BRCA2 antibody. Each point represents the mean of one replicate, normalized to signals in control HeLa cells (siNT -MMC). Shown are Mean ± SEM, data were analysed by a two-way ANOVA, with Tukey’s Multiple Comparison test. (B) Representative PLA images described in (A), showing more BRCA2-PALB2 events (white) in HeLa cells silenced for DPP9 compared to cells treated with non-targeting siRNA (siNT). Actin (Phalloidin) is shown in green and the nucleus (DAPI) in blue. Scale bar 10 µm. (C) Cycloheximide (CHX)-chase assays in HeLa WT cells, showing a rapid turnover of endogenous BRCA2 following exposure to 300 nM MMC (24 hours). Vinculin was analysed as a loading control. (D) Graph summarizing three independent CHX-chase assays performed as described in (C). The ratios of BRCA2 to Vinculin were calculated using 100% at time 0 hours. Mean ± SEM, paired two-tailed t-test. (E) CHX-chase assays in the presence of MMC showing that BRCA2 is more stable in HeLa DPP9 KD cells than in the corresponding HeLa WT cells. (F) Graph summarizing four independent CHX-chase assays performed as shown in (E). Mean ± SEM, paired two-tailed t-test. (G) CHX-chase of HeLa WT cells in the presence of MMC, revealing a greater stability of BRCA2 in DPP9 silenced cells compared to cells treated with non-targeting siRNA oligonucleotides. (H) Graph summarizing three independent CHX-chase assays performed as shown in (G). Mean ± SEM, paired two-tailed t-test. (I) HeLa WT cells were transfected with BRCA2_1-1000_ and BRCA2ΔMP _3-1000_ constructs (both tagged at the carboxy terminus with FLAG) and co-transfected with a GFP expressing plasmid, as transfection and loading control. Graph summarizing three independent CHX-chase assays performed in HeLa WT cells transfected with BRCA2_1-1000_ and BRCA2ΔMP _3-1000_-FLAG constructs. BRCA2-FLAG signals are related to the transfection and loading control GFP. BRCA2 level in relation to GFP were defined as 100 % at time 0 hours. Mean ± SEM, paired two-tailed t-test. (J) Representative Western Blots of transfected cells described in (I). Control cells were transfected with GFP only (U.T).

**Figure 4 - figure supplement 1:**
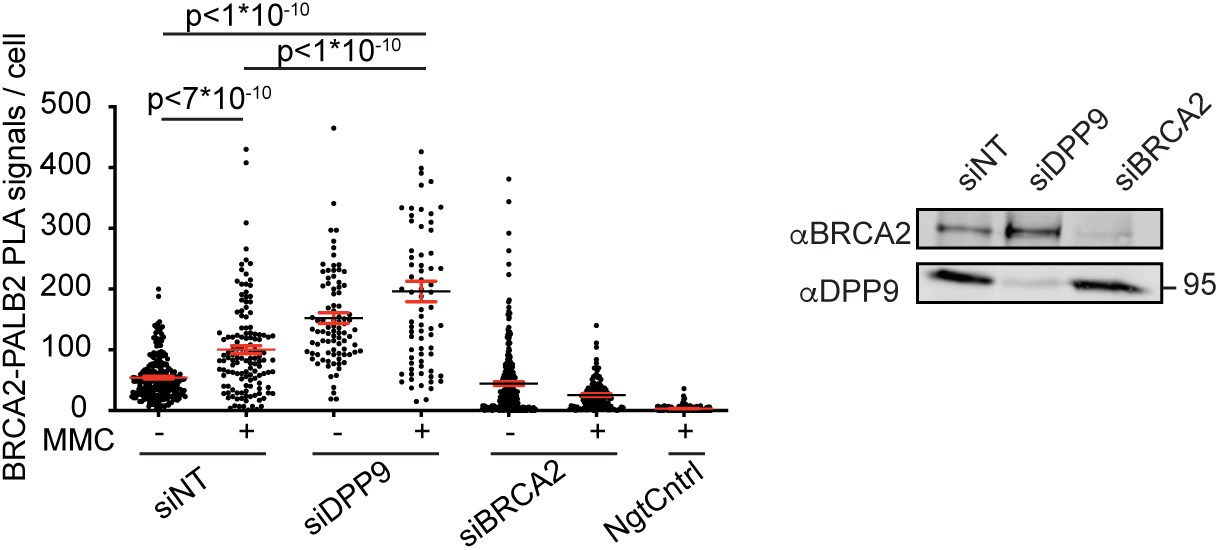
more BRCA2-PALB2 events in HeLa cells silenced for DPP9. PLA quantification showing more BRCA2-PALB2 events in HeLa cells silenced for DPP9 compared to cells treated with non-targeting siRNA (siNT). PLA of the technical control samples (NgtCntrl) did not include the BRCA2 antibody. Actin (Phalloidin) is shown in green and the nucleus (DAPI) in blue. Shown is one representative of 4 independent experiments. Mean ± SEM, data were analysed by a two-way ANOVA with Tukey’s Multiple Comparison test. Western blot showing steady state protein levels of these samples.

**Figure 4 - figure supplement 2:**
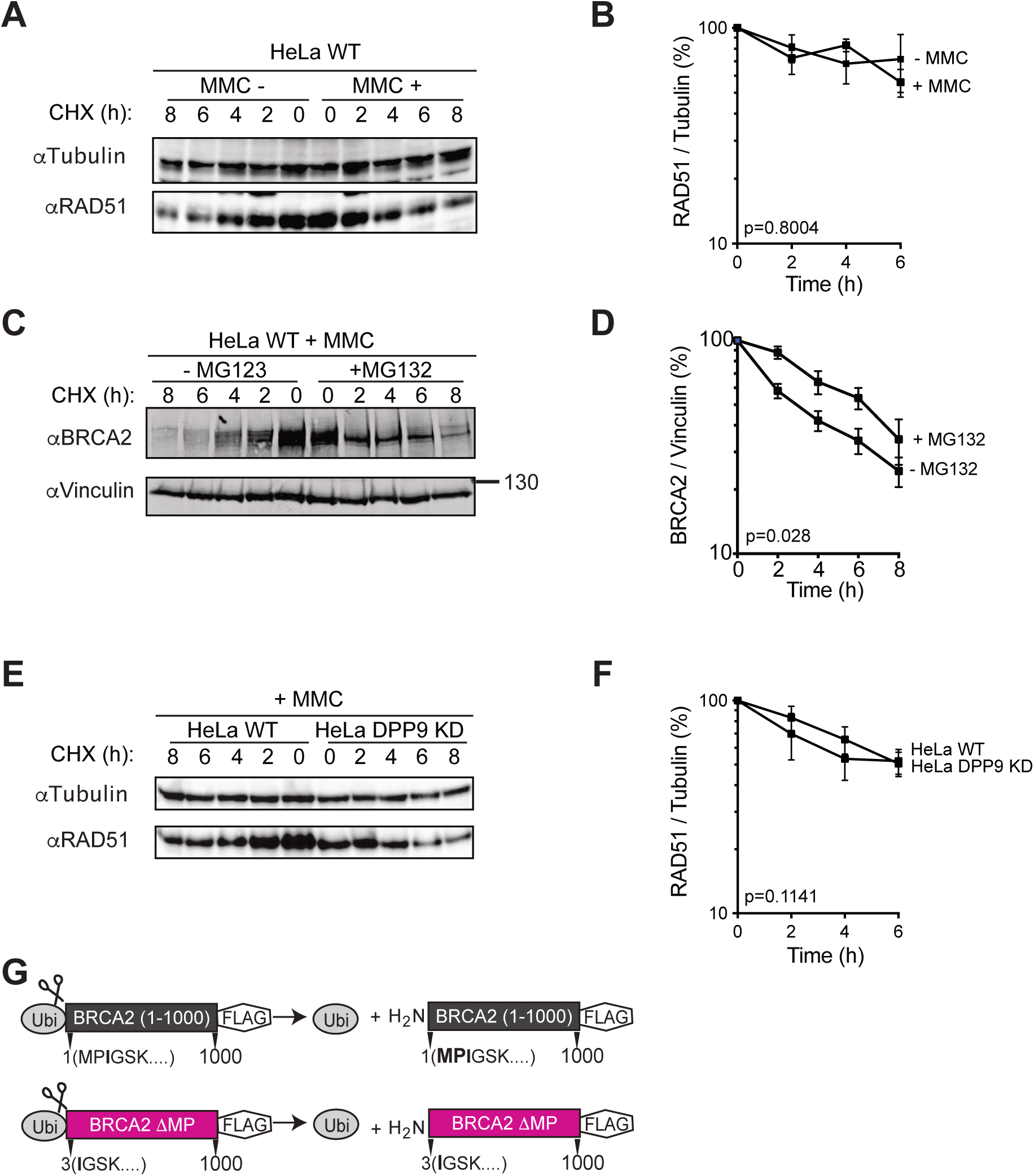
RAD51 turnover is not affected by DPP9. (A) Cycloheximide (CHX)-chase assays in HeLa WT cells, showing that the turnover of endogenous RAD51 is unaltered by exposure to MMC (300 nM, 24 h). Tubulin was analyzed as a loading control. B) Graph summarizing three independent CHX-chase assays performed as described in (A). The ratio of RAD51 to tubulin were defined as 100% at time 0 hours. Mean ± SEM, paired t-test. (C) CHX-chase of HeLa WT cells in the presence of MMC, showing that BRCA2 is more stable in cells treated with the proteasome inhibitor MG132 (100 µM). (D) Graph summarizing three independent CHX-chase assays performed as described in (C). Mean ± SEM, paired t-test. (E) CHX-chase of HeLa WT and HeLa DPP9 KD cells in the presence of MMC, showing that RAD51 turnover if not affected by DPP9. (F) Graph summarizing three independent CHX-chase assays performed as described in (E). Mean ± SEM, paired t-test. (G) Graphical presentation of the plasmids transfected in (E and F).

To address this question, we performed cycloheximide (CHX) chase assays. These demonstrated that endogenous BRCA2 is relatively stable with a half-life greater than 8 hours, and that the addition of MMC stimulates an accelerated turnover of BRCA2, displaying a considerably shorter half-life of 2-3 hours, while the stability of RAD51 was unaltered (Figure 4C and 4D, Figure 4 - figure supplement 2A and 2B). The rapid MMC-induced degradation of BRCA2 was less pronounced in the presence of the proteasome inhibitor MG132, pointing to a role of the ubiquitin-proteasome pathway (Figure 4 - figure supplement 2C and 2D). CHX chase assays were also conducted in the DPP9 KD cells, revealing that while the half-life of RAD51 was not affected, the MMC-triggered degradation of BRCA2 was significantly less pronounced in the HeLa DPP9 KD cells compared to HeLa wild-type cells (Figure 4E and 4F, Figure 4 - figure supplement 2E and 2F). A similar stabilization of BRCA2 was observed in cells silenced for DPP9 (Figure 4G and 4H).

Next we asked whether BRCA2 stability is indeed mediated by the N-terminal dipeptide Met_1_-Pro_2_, which we showed to be removed by DPP9 (Figure 2). To this end, a BRCA2ΔMP_(3-1000)_ mutant lacking these two residues was cloned to mimic a BRCA2 product which would be generated following cleavage by DPP9 (MP↓<u>I</u>GSK → IGSK) (Figure 4 - figure supplement 2G). To produce the required N-terminus (without the initiator methionine) an N-terminal ubiquitin tag was fused to the BRCA2 constructs. This technique relies on endogenous ubiquitin proteases that remove the ubiquitin thereby exposing the desired N-terminus (Varshavsky 2005). CHX chase assays show that the variant containing the unmodified N-terminus BRCA2_(1-1000)_ was relatively stable with a half-life greater than 8 hours, similar to that of the endogenous BRCA2 in the absence of MMC. On the other hand, BRCA2ΔMP_(3-1000)_, was significantly less stable, with a half-life of 2-3 hours (Figure 4I and 4J), similar to the turnover rate of BRCA2 following MMC exposure.

Taken together, we show that DPP9 limits the interaction of BRCA2 with PALB2 and targets BRCA2 for degradation. DPP9-mediated destabilization of BRCA2 is triggered by the presence of DNA-damage, and is mimicked by the BRCA2ΔMP_(3-1000)_ truncation mutant.

### DPP9 activity promotes RAD51 filament formation

Our findings show that although DPP9 triggers the degradation of BRCA2 following DNA damage, DPP9 activity is critical for cell survival under these conditions. The main role of BRCA2 in repair of DSB is to promote the assembly of RAD51 filaments on the ssDNA overhangs (Thorslund et al. 2010; Jensen et al. 2010; Jie Liu et al. 2010; Jasin & Rothstein 2013). Thus, we aimed to better understand these seemingly conflicting observations, by examining the effect of DPP9 on RAD51, the downstream interactor of BRCA2.

Pulse chase experiments have revealed that in contrast to BRCA2, the turnover of RAD51 is not altered in DPP9-deprived cells (Figure 4 - figure supplement 2E and 2F), suggesting that the effect of DPP9 is specific for BRCA2. However, fractionation assays showed that less RAD51 is present in chromatin fractions from HeLa DPP9 KD cells compared to those from HeLa WT cells (Figure 5A). Furthermore, cells silenced for DPP9 did not display RAD51 foci (RAD51 filaments) in response to MMC (Figure 5B and 5C, Figure 5 - figure supplement 1A and 1B). Consistently, fewer RAD51 foci were observed in the gki MEF DPP9^S729A^ cells in response to MMC, in comparison to the MEF WT cells (Figure 5D and 5E, Figure 5 - figure supplement 1C), strongly suggesting that the enzymatic activity of DPP9 is necessary to promote the assembly of RAD51 filaments.

**Figure 5.**
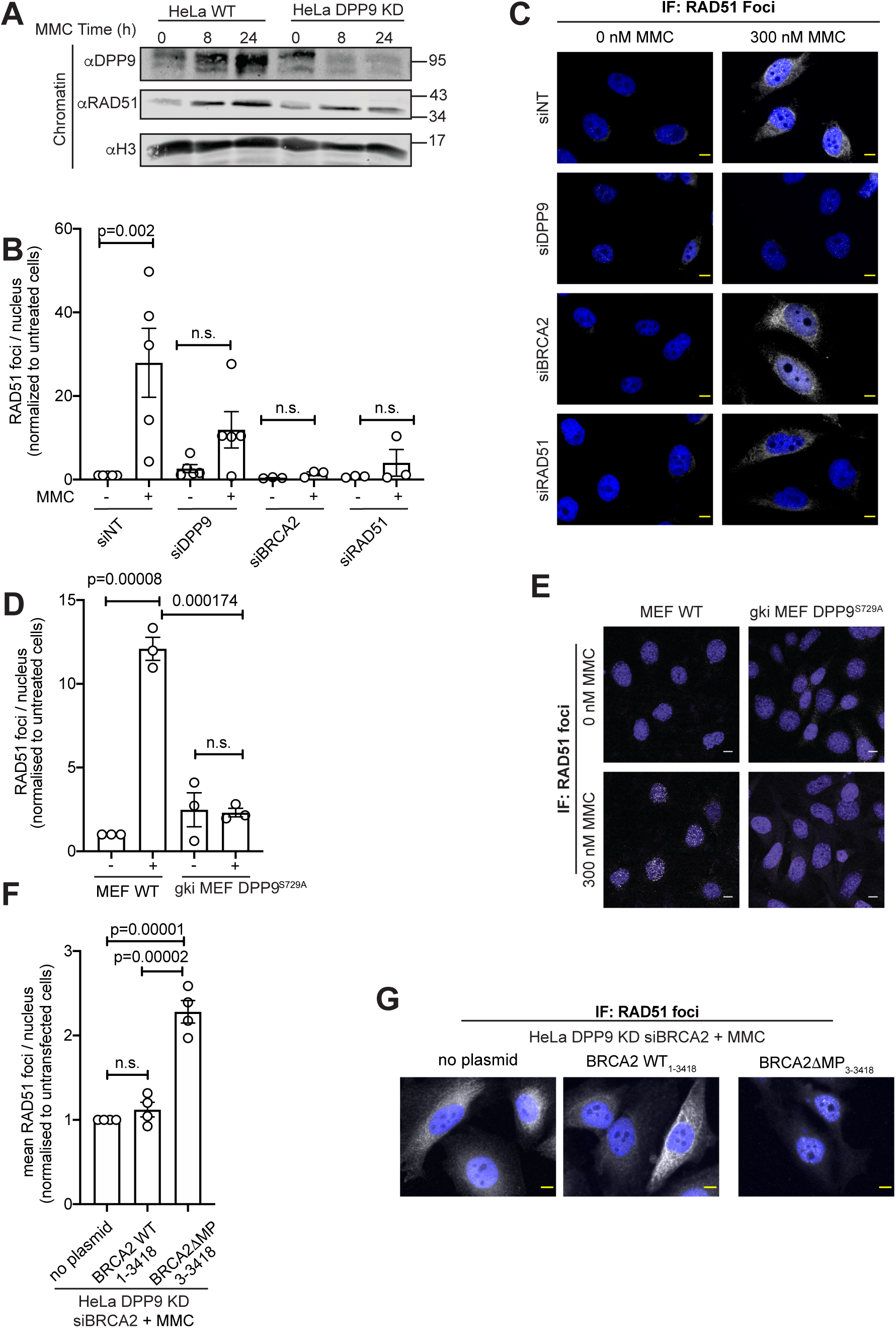
Silencing of DPP9 disrupts RAD51 foci formation. (A) Less RAD51 is associated with chromatin following exposure to MMC in HeLa DPP9 KD cells compared to HeLa WT cells. Histone 3 (H3) was assayed as loading control. Shown is a representative Western Blot of three independent experiments. (B) Graph summarizing four independent biological replicates of RAD51 showing that fewer RAD51 foci are present in response to MMC in cells silenced for DPP9 compared to HeLa cells targeted with non-coding siRNA (siNT). In each replicate, more than 60 cells were quantified for each condition respectively. Each point represents the mean of one biological replicate, normalized to signals in untreated HeLa cells (siNT-MMC). Shown are Mean ± SEM, data were analysed by a two-way ANOVA with Tukey’s Multiple Comparison test. (C) Representative immunofluorescence images of RAD51 as described in (B). Control cells were silenced for BRCA2 or RAD51. The nucleus (DAPI) is shown in blue, RAD51 in white. Scale bar 10 µm. (D) Graph summarizing three independent biological replicates of RAD51 showing fewer MMC-triggered RAD51 foci in gki MEF DPP9^S729A^ compared to MEF WT cells. Each dot represents the mean of one biological replicate, normalized to signals in MEF WT cells – MMC. In each replicate, more than 60 cells were quantified for each condition respectively. Mean ± SEM, data were analysed by a two-way ANOVA with Tukey’s Multiple Comparison test. (E) Immunofluorescence images revealing fewer RAD51 foci (white) in gki MEF DPP9^S729A^ cells expressing inactive DPP9 compared to the MEF WT cells following exposure to MMC. The nucleus (DAPI) is shown in blue, RAD51 in white. Scale bar 10 µm. (F) Significantly more RAD51 foci are present in cells transfected BRCA2ΔMP (3-3418) compared to BRCA2 silenced cells or cells expressing the untruncated BRCA2. HeLa DPP9 KD cells were silenced for BRCA2 and transfected with BRCA2ΔMP_(3-3418)_ or BRCA2 _(1-3418)_ with the untruncated N-terminus. The graph summarises the results from four independent biological replicates. In each replicate, more than 60 cells were quantified for each condition respectively. Each point represents the mean number of RAD51foci of one biological replicate, normalized to signals in HeLa DPP9 KD cells silenced for BRCA2. Shown are Mean ± SEM, data were analysed by a one-way ANOVA. (H) Representative immunofluorescence images of cells described in (G) showing the formation of RAD51 foci in HeLa DPP9 KD cells transfected with full-length BRCA2 (1-3418) or BRCA2ΔMP (3-3418), following exposure to MMC. The nucleus (DAPI) is shown in blue, RAD51 in white. Scale bar 10µm.

**Figure 5 - figure supplement 1:**
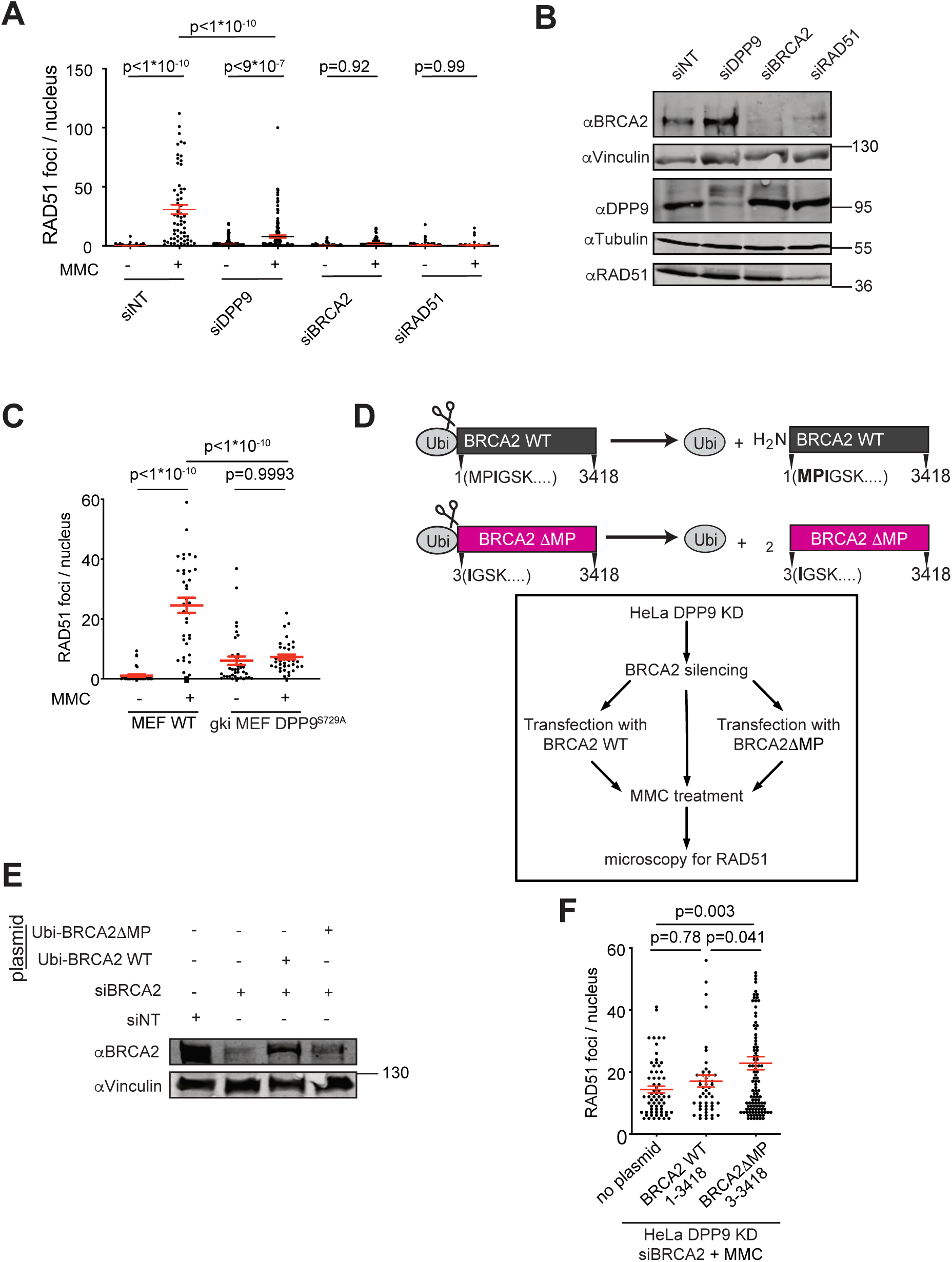
Silencing of DPP9 disrupts RAD51 foci formation. (A) Fewer RAD51 foci are formed in response to MMC in HeLa cells silenced for DPP9, compared to control cells silenced with non-targeting siRNA. Shown is a representative quantification of one biological replicate. Signals of more than 60 cells were quantified for each condition respectively. Mean ± SEM, data were analysed by a two-way ANOVA with Tukey’s Multiple Comparison test. Representative immunofluorescence images of this experiment are shown in Figure 5C. (B) Western Blot showing the protein levels of the corresponding silenced cells used for the immunofluorescence shown in (A). (C) Shown is a representative quantification of one biological replicate showing fewer RAD51 foci in gki MEF DPP9^S729A^ compared to MEF WT cells. Per experiment, signals of more than 50 cells were quantified for each condition respectively. Mean ± SEM, data were analysed by a two-way ANOVA with Tukey’s Multiple Comparison test. (D) Graphical presentation of the two constructs transfected into HeLa DPP9 KD cells, silenced for BRCA2 (E) and (F) and the graphical representation of the silencing and transfection scheme. (E) Western Blot of HeLa DPP9 KD cells that were silenced for endogenous BRCA2 (siBRCA2) and transfected with full-length BRCA2 (1-3418) or with a BRCA2ΔMP (3-3418) construct. Both constructs were fused to an N-terminal ubiquitin tag, which is removed in the cells to expose the desired N-terminus. (F) Significantly more RAD51 foci are present in cells transfected BRCA2ΔMP_(3-3418)_ compared to BRCA2 silenced cells or cells expressing the untruncated BRCA2. Shown is a representative quantification of one biological replicate. HeLa DPP9 KD cells that were silenced for endogenous BRCA2 (siBRCA2) and transfected with full-length BRCA2_(1-3418)_ or with a BRCA2ΔMP_(3-3418)_ construct. Both constructs were fused to an N-terminal ubiquitin tag, which is removed in the cells to expose the desired N-terminus (D)

Finally, we asked whether the reduced formation of RAD51 foci in DPP9-deprived cells is caused by the processing of BRCA2 N-terminus. More specifically we asked what is the role of the dipeptide Met_1_-Pro_2_ in BRCA2 N-terminus. Thus, we compared the capacity of full length BRCA2 (BRCA2_1-3418_) and a BRCA2ΔMP truncation variant (BRCA2ΔMP_3-3418_) to recover the RAD51 foci formation in BRCA2-silenced cells (Figure 5F and – 5G, Figure 5 - figure supplement 1D-F). To prevent the processing of the full length BRCA2 (BRCA2_1-3418_) by DPP9, both constructs were transfected into HeLa DPP9 KD cells. A silent mutation was introduced into both plasmids to acquire resistance to the BRCA2 siRNA. In line with the pulse chase experiments revealing that BRCA2 mutants lacking the N-terminal Met_1_-Pro_2_ are less stable (Figure 4I and J), BRCA2ΔMP_3-3418_ protein levels were lower than that of the WT BRCA2_1-3418_ variant (Figure 5 - figure supplement 1E), although the same amount of plasmid was transfected. Nonetheless, a significant increase in the RAD51 foci was measured in cells expressing BRCA2ΔMP_3-3418_ but not in those expressing WT BRCA2_1-3418_ (Figure 5F and 5G, Figure 5 - figure supplement 1E and 1F). Taken together these results strongly suggest that DPP9 regulates RAD51 filament formation by processing the N-terminus of BRCA2.

## Discussion

Here we report an unexpected role for DPP9 activity in promoting DNA damage repair. We show that DPP9-deprived cells are hypersensitive to genotoxic agents, accumulate higher γH2AX levels, and show fewer RAD51 foci. Moreover, we identify a previously undescribed interaction between the tumour suppressor BRCA2 and DPP9, which in cells requires access to DPP9 active site. Mass spectrometry analysis shows cleavage and removal of the N-terminal dipeptide Met_1_-Pro_2_ from the BRCA2_1-40_ peptide. Moreover, crystallographic structures of DPP9 crystals soaked with the BRCA2_1-40_ peptide, reveal the dipeptide Met_1_-Pro_2_ of BRCA2 captured in the active site of DPP9, demonstrating intra-crystalline enzymatic activity.

### DPP9 targets BRCA2 to the N-degron pathway

Functionally, DPP9 determines the turnover of BRCA2. Regulated protein degradation is an essential response for maintenance of cellular homeostasis. It is important for removal of mutated or damaged proteins and to establish a correct protein stoichiometry. In this context, BRCA2 undergoes degradation in response to mild hyperthermia (Krawczyk et al. 2011), and upon silencing of the BRCA2-binding protein DSS1 (Li et al. 2006), conditions which could lead to protein aggregation.

BRCA2 protein levels are also known to be lower following DNA damage (Schoenfeld et al. 2004; Jinping Liu et al. 2017). CHX chase assays presented here confirm that DNA damage triggers an accelerated turnover of BRCA2, a process requiring the proteasome. More importantly, we show that DPP9, which cleaves only two residues from the N-termini of its substrates, initiates the MMC-mediated degradation of BRCA2, a protein of 384 kDa. Furthermore, this accelerated turnover is mimicked by a BRCA2 truncation mutant lacking the N-terminal Met_1_-Pro_2_ dipeptide.

The degradation of proteins based on the sequence of their N-terminus is known as the N-degron pathway, and in eukaryotes requires the ubiquitin-proteasome pathway (Varshavsky 2019). Previously we showed that by trimming the N-terminus of Syk (Met-Ala), DPP9 targets the kinase to the N-degron pathway (Justa-Schuch et al. 2016). The results presented here show that similarly, the removal of the N-terminal Met_1_-Pro_2_ dipeptide, converts BRCA2 to an unstable protein, demonstrating BRCA2 to be an N-degron substrate. Notably, N-terminal proline residues are characteristic of extremely poor N-degron substrates (Bachmair et al. 1986). An exception are gluconeogenic enzymes with N-terminal prolines that are highly stable, but targeted for the N-degron pathway by a class of E3 ligases that specifically ubiquitinate these enzymes upon transition of yeast to a glucose-containing medium (S.-J. Chen et al. 2017; Dong et al. 2018; Melnykov et al. 2019). Here, we add DPP9 as an alternative route that cells apply to target proteins with a stabilizing N-terminal proline, by converting these to N-degrons. Similar to the induced degradation of the gluconeogenic enzymes by the N-degron pathway, also the degradation of BRCA2 is not constitutive, but instead triggered by DNA damage.

### DPP9 activity regulates RAD51 foci formation

Here we show that although it accelerates the turnover of BRCA2 in response to DNA damage, DPP9 promotes RAD51 foci formation, a process mimicked by the BRCA2ΔMP mutants. These observations suggest that cleavage of BRCA2 N-terminus by DPP9 may have a dual function: activation of BRCA2 in addition to BRCA2 degradation. While we cannot exclude that BRCA2ΔMP is more active compared to BRCA2 with the unprocessed N-terminus, this scenario is not highly likely since the BRC repeats which mediate the interactions of BRCA2 with several RAD51 copies are located to the centre of BRCA2, and an additional RAD51 binding domain locates to the C-terminus (A. A. Davies et al. 2001; Pellegrini et al. 2002; Esashi et al. 2007; O. R. Davies & Pellegrini 2007).

An alternative possible explanation is that the DPP9-mediated degradation of BRCA2 may positively regulate DNA repair. Turnover of key DNA repair proteins is emerging as a general concept. For example, the SUMO-targeted ubiquitin E3 ligase RNF4 is required for DSBs repair by targeting MDC1 and RPA1 at DNA-damage sites (Galanty et al. 2012; Luo et al. 2012; Yin et al. 2012). Similarly, the ubiquitin E3 ligase RFWD3, facilitates the HR repair by ubiquitinating RAD51 and RPA, leading to the removal of both proteins from DSB sites (Gong & J. Chen 2011; Shangfeng Liu et al. 2011; Elia et al. 2015; Inano et al. 2017; Feeney et al. 2017). However, the majority of the BRCA2-DPP9 PLA events are cytosolic, suggesting that the DPP9-mediated turnover of BRCA2 may play a role other than its removal from the DSBs. Several publications have previously shown that BRCA2 promotes RAD51 filament assembly by binding simultaneously to several molecules of RAD51 and that maintenance of the sub-stoichiometric ratio of BRCA2 compared to that of RAD51 is critical for filament formation (Yang et al. 2005; Thorslund et al. 2010; Jie Liu et al. 2010; Jensen et al. 2010; Shahid et al. 2014). The importance of a lower BRCA2 concentration relative to that of RAD51 was also documented in cellular assays (Magwood et al. 2012). Thus, DPP9-mediated degradation of BRCA2 following DNA damage, may promote RAD51 filament formation by reducing the relative cellular concentration of BRCA2, while those of RAD51 remain unaltered (Figure 6). Of note, similar to DPP9, also USP21 acts as a positive mediator of BRCA2. In contrast to DPP9, USP21 deubiquitinates BRCA2 to increase BRCA2 stability (Jinping Liu et al. 2017). Taken together with the data presented here, we suggest that BRCA2 steady state levels are tightly regulated and respond quickly to changing cellular conditions. Thus, the amount of BRCA2 in response to DNA-damage is determined by DPP9, but can be fine-tuned through the down-stream deubiquitinating enzyme USP21. How DNA damage signals for DPP9 to process BRCA2 will be addressed in future work.

**Figure 6.**
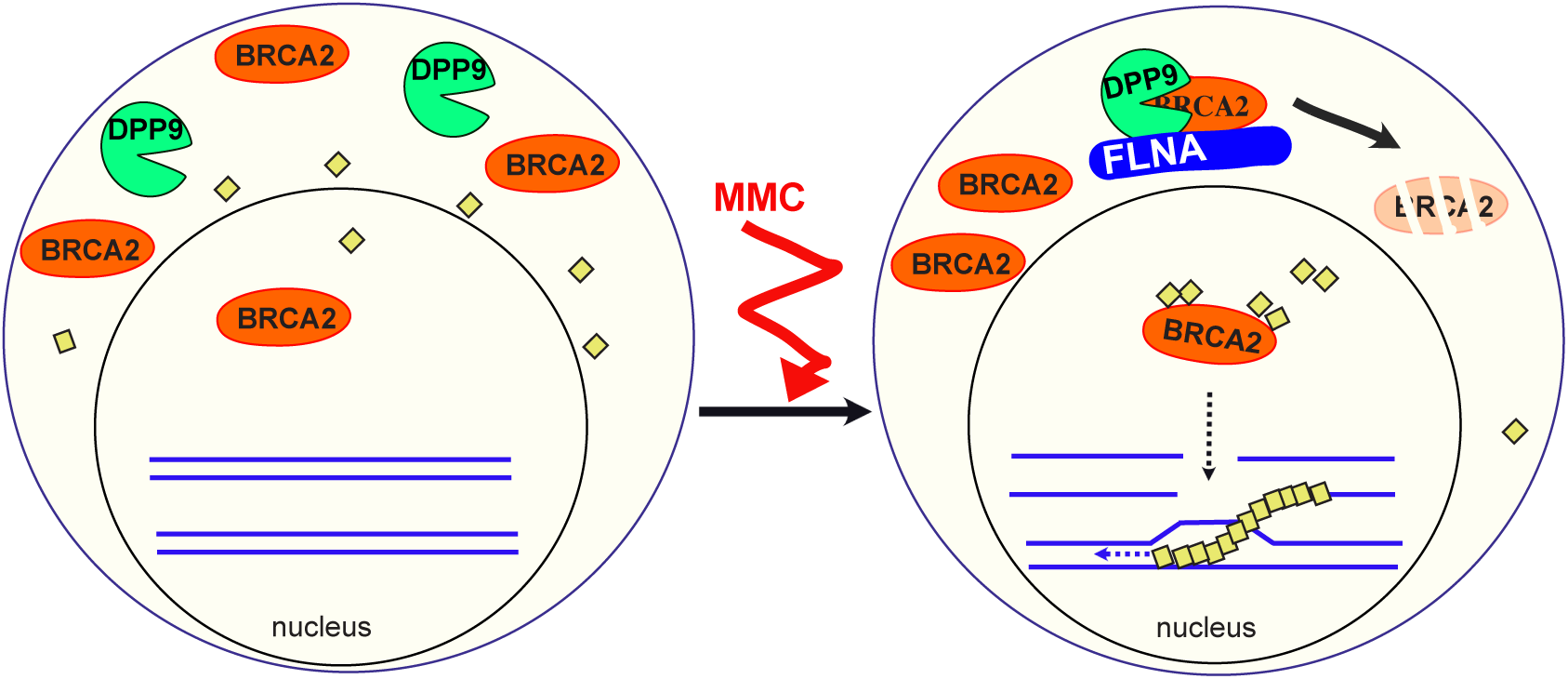
Model: DPP9 promotes DNA damage repair by triggering the degradation of BRCA2. MMC triggers the trimming of BRCA2 N-terminus by DPP9 resulting in its degradation by the N-degron pathway, while RAD51 remains stable. This reduction of BRCA2 protein levels changes the ratio of RAD51 to BRCA2. A high RAD51 and lower BRCA2 stoichiometry has been shown to be beneficial for the formation of RAD51 filaments.

**Figure 6 - figure supplement 1:**
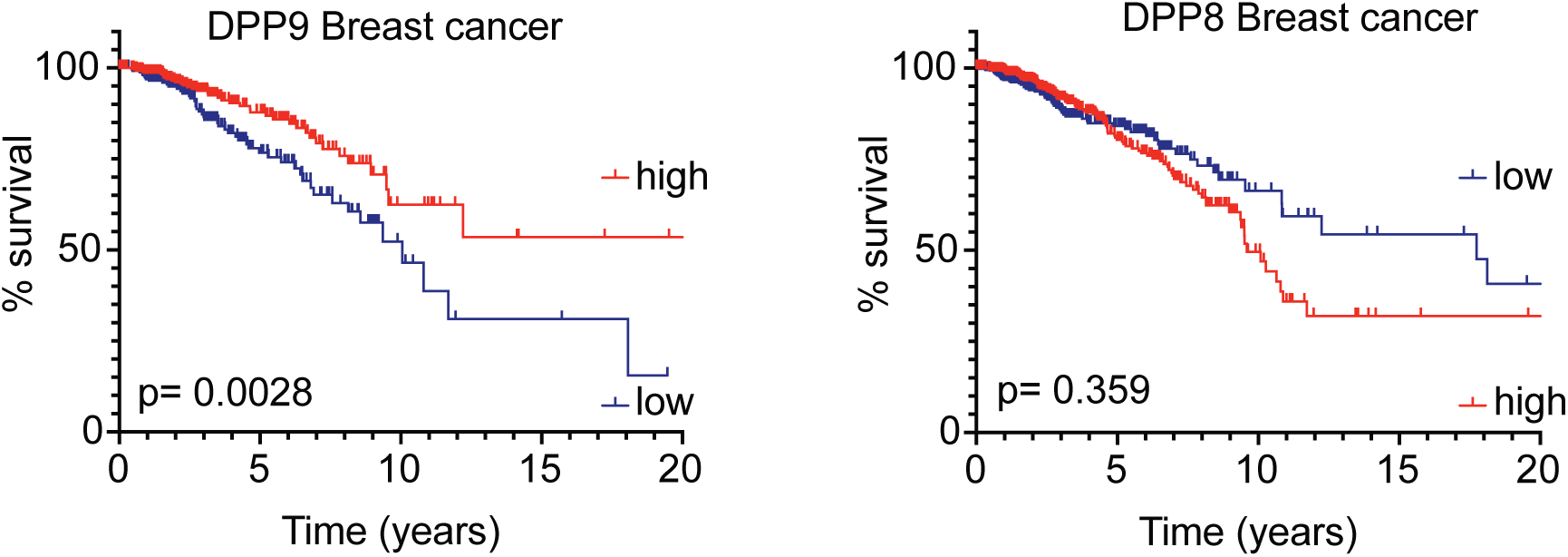
Low DPP9 mRNA expression correlate with poor overall survival for patients with breast cancer. Graphs show separation of Breast cancer patients from The Human Protein Atlas into “high DPP9 or DPP8” (n=282, 445) and “low DPP9 or DPP8” (n =408, 630) mRNA expression (greater than or less than 9.678 or 4.36 reads per kilobase per million, respectively). Results are displayed in Kaplan-Meier survival plots; p values were calculated by log-rank (Mantel- Cox). These revealed a better overall survival in patients with higher levels of DPP9 expression, while differences in the DPP8 levels were not statistically significant.

### Outlook

Carriers of inherited mutations in BRCA2 have a high risk (> 50 %) to develop breast cancer (Mavaddat et al. 2013). Since DPP9 regulates BRCA2, we postulate that deregulation of DPP9 may lead to accumulation of unrepaired DSBs which could lead to cancer development. In this respect, data from The Human Protein Atlas show a correlation between low DPP9 mRNA expression and poor overall survival for patients with breast cancer compared to individuals with tumors displaying higher DPP9 expression who showed a better overall survival (Figure 6 - figure supplement 1) (Human Protein Atlas, (Justa-Schuch et al. 2016)). Such a correlation is not observed for its close homolog DPP8.

Interestingly, DPP9 fusion genes which lead to a premature stop codon have been identified in patients with a high-grade serous ovarian carcinoma, probably leading to lower DPP9 protein expression (Smebye et al. 2017). Such DPP9-fusions mimic the situation in DPP9 KD and gki DPP9^S729A^ MEF cells thus, we predict that defects in BRCA2 interactions and degradation in these patients, resulting in accumulation of unrepaired DSBs. Finally, cells expressing lower DPP9 levels display a high sensitivity to MMC, γ radiation and Olaparib. Thus, DPP9 inhibition in combination with for example Olaparib or radiation, may represent future potential therapies for breast-cancer.

## Materials and Methods

### Key Resources Tables

**Table 2:** Antibodies.

| Antibodies | Method | Source | Identifier |
| --- | --- | --- | --- |
| BRCA2 Mouse | IF/PLA | R&D | MAB2476 |
| BRCA2 Rabbit | IF/PLA | Santa Cruz | sc-8326 |
| BRCA 2 Rabbit | WB | Abcam | 123491 |
| DPP9 Rabbit | WB | Abcam | AB42080 |
| DPP9 Goat | IF/PLA | Self-made | (Justa-Schuch et al. 2016) |
| Filamin A Rabbit | WB | Novus | nb-100-58812 |
| Filamin A Mouse | IF/PLA | Millipore | MAB1680 |
| H2AX-P Rabbit | WB | Cell Signaling | #9718 |
| H2AX-P Mouse | IF | Millipore | #05-636 |
| H3 Rabbit | WB | Cell Signaling | #9715 |
| PALB2 Rabbit | IF/PLA | Bethyl/biomol | A301-246M |
| Phalloidin 488 | IF/PLA | Abcam | Ab176753 |
| RAD51 Rabbit | IF/WB | Abcam | ab63801 |
| Tubulin Mouse | WB | Santa Cruz | sc-32293 |
| Vinculin Mouse | WB | Sigma | V9131 |

**Table 3:** Chemicals.

| Chemical | Source | Identifier |
| --- | --- | --- |
| 1G244 (in DMSO) | AK Scientific, Inc. | #Y0432 |
| CellTiter-Glo® Luminescent Cell Viability Assay | Promega | #G7571 |
| Cycloheximide (CHX) | Sigma | #C7698 |
| Lipofectamine RNAimax | Thermo Fisher | #13778150 |
| Mayer's Hemalum | Merck | #1.09249.0500 |
| Mitomycin C (MMC) | Sigma | #M4287 |
| MG132 (in DMSO) | Sigma | #C2211 |
| Neocarzinostatin (NCS) | Sigma | #N9162 |
| Olaparib | Selleck Chemicals | #AZD2281 |
| Puromycin | Sigma | #P8833 |
| Aprotinin | Carl ROTH | A162.3 |
| Leupeptin | Carl ROTH | CN33.4 |
| Pepstatin | Carl ROTH | 2936.3 |
| Pierce Anti-HA magnetic beads | Thermo Scientific | #88836 |
| Ni-NTA Agarose | Qiagen | #1018244 |
| DuoLink in Situ PLA probe Mouse plus | Sigma | #DUO92001 |
| DuoLink in Situ PLA probe Rabbit plus | Sigma | #DUO92002 |
| DuoLink in Situ PLA probe Mouse minus | Sigma | #DUO92004 |
| DuoLink in Situ PLA probe Goat minus | Sigma | #DUO92006 |
| DuoLink in Situ Detection Reagents Red | Sigma | #DUO92008 |
| Phusion DNA polymerase | NEB | #M0363S |

**Table 4:**
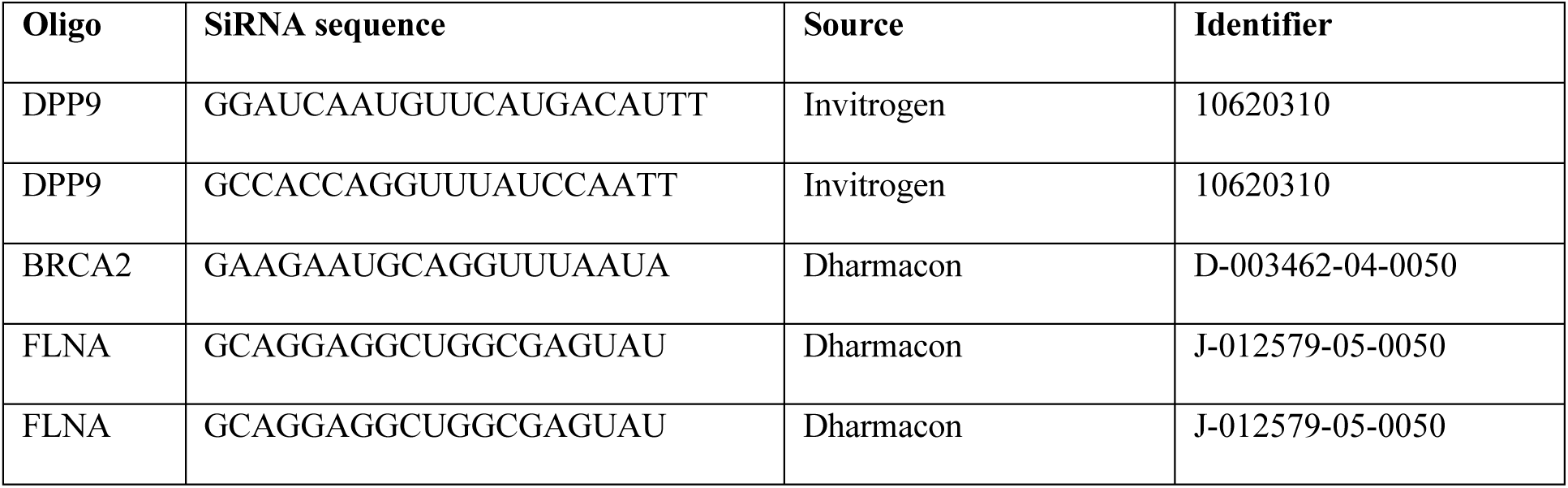
siRNA oligonucleotides.

**Table 5:**
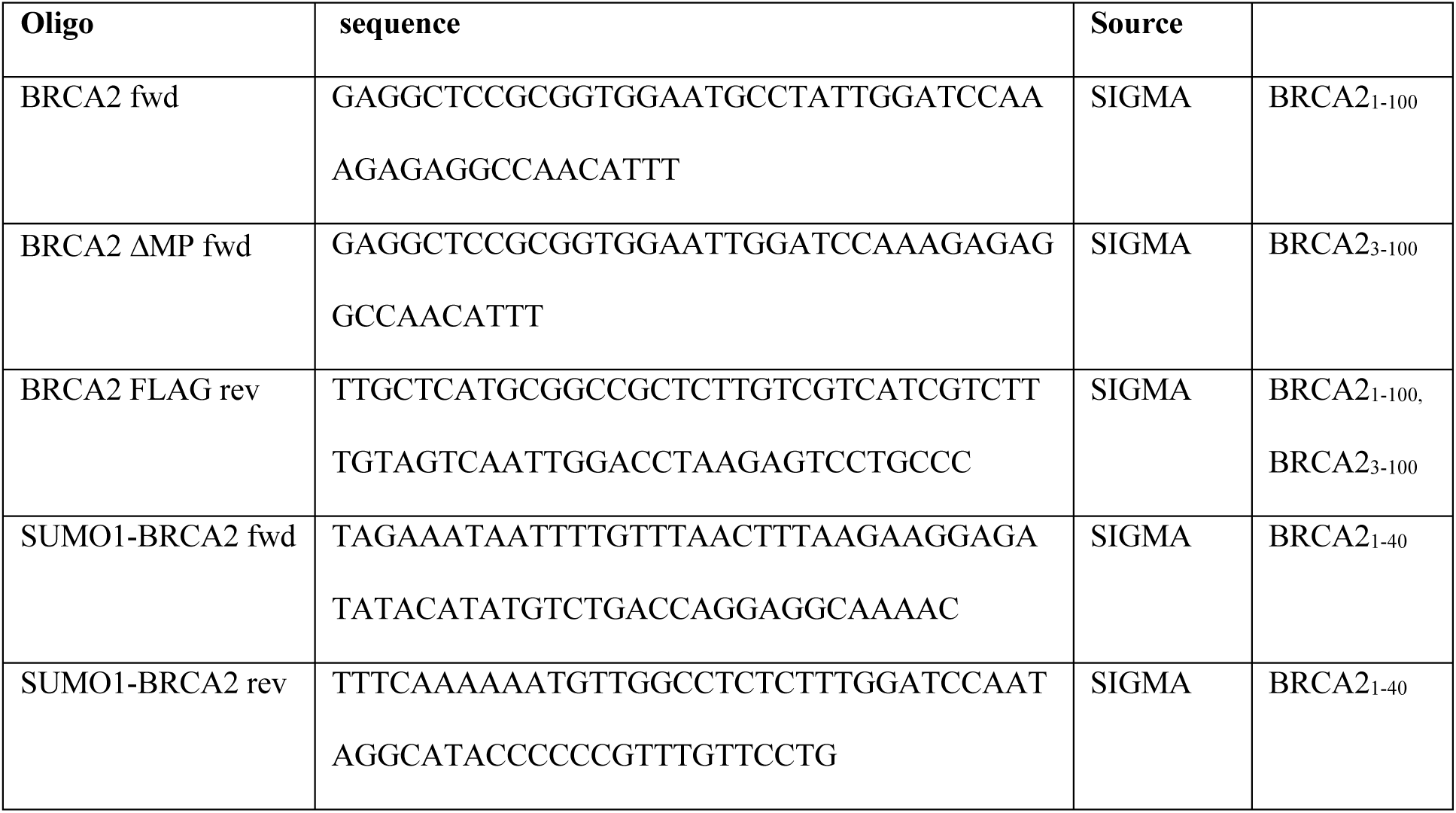
Primers.

**Table 6:**
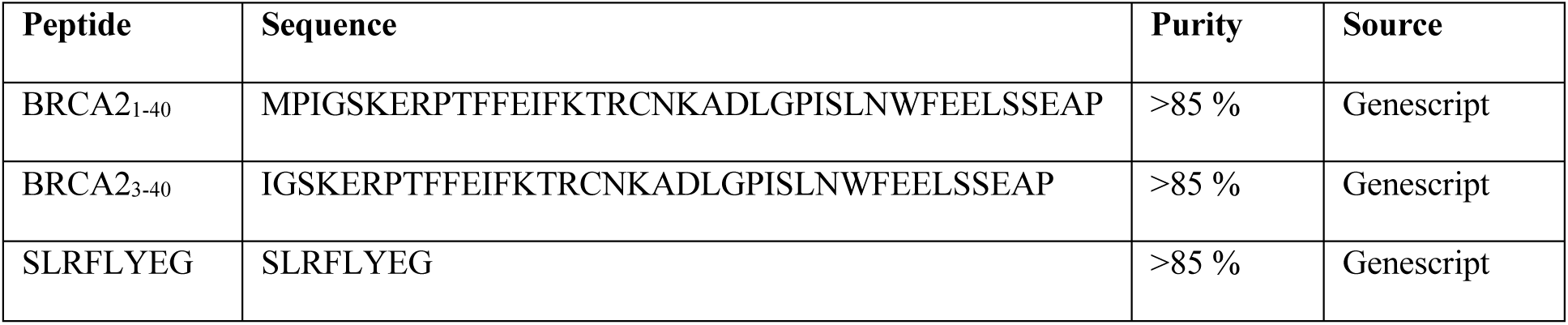
Peptides.

**Table 7:** Software.

| Software | Source |
| --- | --- |
| DuoLink software | Sigma |
| ImageStudio/ ImageStudio Lite 4.0.21 | LI-COR |
| LSM 510 Release Version 4.0 SP2 | Zeiss |
| LSM Image Browser | Zeiss |
| Celigo Software | Celigo |
| NanoDrop 2000 Software | ThermoScientific |
| Prism 8 | Graphpad |

## METHOD DETAILS

### Cell culture

HeLa DPP9 stable knock down cells (DPP9 KD) and the corresponding HeLa WT cells (Genescript) (Justa-Schuch et al. 2016) were cultured at 37°C and 5 % CO2 in Dulbecco’s modified Eagle’s medium supplemented with 10 % fetal bovine serum, 2 mM L-glutamine, 100 U/ml penicillin and 100 μg/ml streptomycin. To maintain the selection pressure on the DPP9 KD cells, 1.5 µg/ml puromycin (Sigma-Aldrich, Germany) was added to the growth medium. MEF wt and gkiMEF DPP9^S729A^ (Gall et al. 2013) cells were cultured at 37 °C and 5% CO2 in Dulbecco’s modified Eagle’s medium supplemented with 10% fetal bovine serum, 2 mM L-glutamine, 100 U/ml penicillin and 100 μg/ml streptomycin.

### Cloning of the BRCA2 fusion vectors

BRCA2 construct covering the first 1000 amino acids (BRCA2_1-1000_) and a corresponding BRCA2 lacking the N-terminal dipeptide Met-Pro (BRCA2_3-1000_) were cloned into pcDNA3.1+ vector with a Ubiquitin tag fused at the N-terminus to provide a defined N-terminus, as first described by Bachmair and Varshavsky (Bachmair et al. 1986). As template we used pcDNA3 236HSC WT BRCA2, a kind gift from Mien-Chie Hung (Wang et al. 2002) obtained from Addgene. SUMO1-BRCA2_1-40_-HA-His constructs were cloned by adding SUMO1 to the N-terminus of BRCA2_1-39_-3HA-His pET11a (custom-made from Genescript), using the Gibson Chew Back and Anneal Assembly (CBA) as described in (Torella et al. 2014). Primers are listed in Table 4. The presence of the inserts was verified by restriction, followed by sequencing.

### Silencing of HeLa cells

For silencing of DPP9, HeLa cells were transfected with lipofectamine RNAi max (Cat #13778150) following the manufacturer’s instructions. Cells were analysed 72 hours after transfection. For MMC treatment, cells were treated with MMC 48 h after silencing and harvested 24 hours later with Laemmli Sample Buffer (0,1 M Tris-HCl pH 6.8, 4% SDS, 20 % glycerol, 0.02% bromophenol blue, 100 mM DTT). Alternatively, cells were fixed on coverslips with 4 % formaldehyde as described below.

### Protein purification

Human recombinant DPP9 was expressed from Sf9 cells and purified essentially as described in (Pilla et al. 2012). For purification of N-terminally His-HA-tagged fragments corresponding to the N-terminus of BRCA2 (BRCA2 _1-40_HA-His), SUMO1BRCA2_1-40_HA-His was transformed into BL21 (DE3 codon+) cells and induced by 1 mM IPTG for 2 h at 30 °C. Cells were lysed in 25 ml Lysis Buffer (50 mM NaH_2_PO_4_,300 mM NaCl, 10 mM Imidazol pH 8.0, 1 mM ß-Mercaptoethanol and protease inhibitors: 1 µg/ml Aprotinin, 1 µg/ml Leupeptin and 1 µg/ml Pepstatin), by sonification. The 100,000 g supernatant was bound to Ni-NTA Agarose (Qiagen) for 2 h. Beads were thoroughly washed with Lysis buffer lacking protease inhibitors and incubated with 20 µg SENP over night to release SUMO1. Following extensive washing with Lysis buffer, BRCA2_1-40_HA-His was eluted with Elution buffer (50 mM Na-Phosphat pH 8,0 300 mM NaCl, 250 mM Imidazol, 1 mM ß-Mercaptoethanol).

### Pull-down assay

To test for an interaction between recombinant DPP9 and a BRCA2 N-terminal fragment, BRCA2 _1-40_HA-His (2 µg) was incubated with magnetic anti-HA beads (Pierce Anti-HA magnetic beads, Cat# 88836, Thermo Scientific) in a Binding buffer containing (25mM TRIS pH 7.5, 150 mM NaCl, 1 mM EDTA, 0.01 % (v/v) IGEPAL, 1 mM DTT), for 1 hour at 4 °C. Next recombinant DPP9 (2 µg) was added and incubated over-night at 4°C. Where stated, DPP9 was preincubated for 1 hour with the following inhibitors: 100 µM 1G244 or 110 µM SLRFYEG. Following extensive washing, with the same buffer, bound proteins were eluted with the HA peptides and analysed by Western Blot.

### Surface plasmon resonance (SPR)

Direct interactions between recombinant DPP9 and BRCA2-derived N-terminal peptides (1-40 and 3-40 amino acids) were analysed employing surface plasmon resonance (SPR). The surface of a NiHC1000m sensorchip (Xantec Bioanalytics, Duesseldorf, Germany) was rinsed with 0.5 M EDTA (pH 8.0) solution and afterwards conditioned in SPR running buffer (20 mM HEPES pH 7.4, 150 mM NaCl, 50 µM EDTA, 0.005 % TWEEN-20). The chip-surface was loaded with Ni^2+^-ions, injecting 300 mM NiSO_4_ solution. A 200 nM solution of His-tagged DPP9 (ligand) was injected over the chip surface and immobilized to a surface density of 800-1000µRIU (Refractive index units; response observed), at a flow rate of 15 µl/min. A serial dilution of the analyte (BRCA2 peptides) diluted in SPR running buffer was injected, and association was followed for 4.5 min. Dissociation of the analyte was monitored for 7 minutes. For each analyte concentration, the surface was regenerated, injecting 0.5 M EDTA pH 8.0 and fresh ligand was immobilized as described above (one immobilization/regeneration cycle for each analyte concentration). All binding experiments were performed on a Reichert SR 7500DC biosensor at 20°C and a flow rate of 40 µl/min. Equilibrium binding analysis was performed using Graph Pad Prism 6.0.

### Peptidase activity assay by liquid chromatography-tandem mass spectrometry (LC/MS/MS)

The assay was essentially performed as described in (Justa-Schuch et al. 2016). 50 µM of the BRCA2 _1-40_ peptide was incubated alone or in the presence of 125 nM DPP9. To test for inhibition, 10 µM peptide inhibitor (SLRFLYEG) was added. Reactions were stopped after 6 hours by dilution and acidification in aqueous 0.1% formic acid, 2% acetonitrile (1/20.000, v:v). In addition, bovine insulin was added to the dilution buffer at a concentration of 1 pmol/ul to prevent analyte loss due to adsorption. The resulting samples were analysed on a nanoLC425 nanoflow chromatography system coupled to a TripleToF 5600+ Plus mass spectrometer of QqToF geometry (both AB SCIEX). 5 µl of sample were pre-concentrated on a self-packed Reversed Phase-C18 precolumn (Reprosil C18-AQ, Pore Size 120 Å, Particle Size 5 µm, 4 cm length, 0.15 cm I.D., Dr. Maisch) and separated on a self-packed Reversed Phase-C18 microcolumn (Reprosil C18-AQ, 120 Å, 3 µm, 15 cm, 0.075 cm) using a 15 minutes linear gradient (5 to 50% acetonitrile, 0.1% formic acid modifier, flow rate 300 nl/min, column temperature 50°C) followed by a 5 minutes high organic cleaning step and a 15 minutes column re-equilibration. The eluent was introduced to the mass spectrometer using a Nanospray III ion source with Desolvation Chamber Interface (AB SCIEX) via a commercial Fused Silica tip (FS360-20-10-N-C15, New Objective) at a spray voltage of 2.2 kV, a sheath gas setting of 15 and an interface heater temperature of 150°C. The MS acquisition cycle consisted of a 500 ms TOF MS survey scan that was used for profiling of substrate and product concentrations followed by data-dependent triggering of up to 5 100 ms TOF product ion spectra to confirm the identity of detected analytes. Data analysis was performed using the Analyst TF 1.7 and the PeakView 2.1 softwares (AB SCIEX). Analyses were performed in quadruplicate.

### Protein crystallography

Purification, crystallization and structure solution of DPP8 and DPP9 was performed as published in our previous report (Ross et al. 2018). Briefly, the DPP8 isoform 1 (Uni-ProtKB Q6V1×1) and DPP9 isoform 2 (Uni-ProtKB Q86TI2-2) were expressed in *Spodoptera frugiperda* cells and purified. The crystallization was performed in both cases using the hanging drop method with 0.46 M Na-citrate pH 6.75 as precipitant solution and 10 mg/ml of DPP8 at 4 °C. DPP9 crystallization occurred at 20 °C, using 20 mg/ml of protein and 10% PEG 8000, 25% glycerol, 0.16 M Calcium acetate, and 0.08 M cacodylate pH 6.25 as precipitant solution. Crystals of space group C222_1_ (DPP8) and P12_1_1 (DPP9) were soaked with BRAC2_1-40_ peptide performing dropping using a pico-dropper (Ross et al. 2018). Data were collected at SLS-X10SA and solved to 3.2 Å and 3.0 Å, respectively, by molecular replacement using a DPP8 structure (PDB: 6EOO). Data processing statistics and refinement values are summarized in table 1. PDB: 6QZW shows DPP8 bound to a dipeptide (MP) from the N-terminus of BRCA2

PDB: 6QZV shows DPP9 bound to a dipeptide (MP) from the N-terminus of BRCA2

### Cycloheximide (CHX) chase assays

HeLa WT cells were seeded at 0.5×10^6^ density in 6 well plates and grown for 24 hours. Next, 300 nM MMC was added to the cells following an additional 24 h incubation. The CHX chase was started by the addition of 100 μg/ml CHX. Where stated, 100 μM of MG132 was added to the cells 30 min before addition of the CHX. Samples were harvested at the indicated time points directly in Laemmli Sample Buffer. Before loading on SDS PAGE, samples were incubated for 10 min at 65°C followed by 4 pulses of sonication at 1% intensity to allow better detection of BRCA2. Immunoblotting and incubation with primary antibodies were performed according to standard protocols. Secondary fluorophore-coupled antibodies were applied. Signals were developed in the Odyssey Sa infrared Imaging System (LI-COR) and analysed with the ImageStudio (version 4.0.21, LI-COR) software.

### Chromatin fractionation

HeLa WT and HeLa DPP9 KD were seeded in 6 well plates. The next day (90% confluence), cells were treated with MMC (300nM) for 0, 8 or 24 hours. Chromatin fractionation was performed essentially as described in (Kari et al. 2016). Briefly cells were re-suspended in lysis buffer (10 mM HEPES pH 7.9, 10 mM KCl, 1.5 mM MgCl_2_, 0.34 M sucrose, 10% glycerol, 0.1% Triton X-100, 1 mM DTT, and protease inhibitors) and centrifuged at 1500 *g* for 5 minutes. The nuclear pellet was lysed in nuclear lysis buffer (3 mM EDTA, 0.2 mM EGTA, 1 mM DTT, and protease inhibitors) for 30 minutes on ice. Soluble chromatin fractions were separated by centrifuging at 1700 *g* or 5 minutes. Chromatin fractions were sonicated in water sonicator (BioRad) for 15 min before loading on SDS–PAGE electrophoresis. Immunoblotting and antibody incubations were performed as described above.

### Immunofluorescence

HeLa and MEF cells were grown on coverslips in 24 well plates, fixed with 4% formaldehyde in PBS and permeabilized with 0.2% Triton-X-100 in PBS for 5 minutes. Cells were washed with PBS and blocked with 2% BSA in PBS for 10 minutes. Both cell lines were incubated with primary antibodies for 90 minutes at 37°C. Following a PBS wash, cells were incubated for 45 minutes at room temperature with the respective secondary antibodies. Cells were washed with PBS and water and mounted in fluorescent mounting medium (DAKO) with DAPI. Control samples were treated with secondary antibodies only, to estimate background staining. Cells were analysed and images were taken using either an LSM 510-Meta confocal microscope, oil immersion objective 63x/1.3 (Carl Zeiss MicroImaging, Inc), or a Nikon Eclipse Ti2-E Inverted microscope, Plan Apoλ oil immersion objective 60x NA1.40 WD=0.13 (Nikon Instruments Inc). Images were processed using LSM image Browser (Carl Zeiss MicroImaging, Inc) or NIS-Elements AR 5.02.00 (Nikon Instruments Inc), based on the microscope used, and FIJI for the preparation of figures. RAD51 foci were quantified using the Duolink ImageTool (SIGMA-Aldrich).

### Proximity Ligation Assay

Proximity Ligation Assay (PLA) was performed using the DUOLINK In Situ PLA Kit from Sigma Aldrich, according to the manufacturer’s protocol. In short, HeLa cells were grown on coverslips and fixed as described above for the Immunofluorescence assays. In the case of DPP9 inhibition studies, 10 µM 1G244 were added for 30 minutes to the cells before fixation. Control cells were mock treated with DMSO. HeLa cells were then incubated with primary antibodies, 90 minutes at 37°C and actin filaments were simultaneously counterstained with CytoPainter Phalloidin-iFluor 488 Reagent (Abcam - #ab176753). Control slides were treated with only one primary antibody to estimate background staining in each experiment. Next, coverslips were washed with PBS and treated with the PLA reagents. Cells were mounted in DAKO with DAPI fluorescent mounting medium and analysed using a LSM 510-Meta confocal microscope, oil immersion objective 63x/1.3 (Carl Zeiss MicroImaging, Inc) or a Nikon Eclipse Ti2-E Inverted microscope, Plan Apoλ oil immersion objective 60x NA1.40 WD=0.13 (Nikon Instruments Inc). Images were processed using LSM image Browser (Carl Zeiss MicroImaging, Inc) or NIS-Elements AR 5.02.00 (Nikon Instruments Inc), based on the microscope used and subsequently analysed using the Duolink ImageTool (SIGMA).

### Viability assay

Cells were seeded in 96-well White Microplates (Perkin Elmer cat# 6005680) and treated with different concentrations of Olaparib for 96 hours. For the MMC assays, cells were first incubated for 18-24 h to allow cell attachment. Next, different MMC concentrations were added and cells were analysed 72 hours later. Control cells were treated with DMSO. Cell viability was measured using the CellTiter-Glo®Luminescence Cell Viability Assay (Promega, cat# Cat G7571). CellTiter-Glo®Reagent was added in a 1:1 ratio to the cell culture medium in a well. The plate was placed on an orbital shaker for 10 minutes for induction of cell lysis. Subsequently, the luciferase signal was measured on a LuminometerDLReadyTMCentro LB 960 reader. Each experiment was performed three times, in triplicate.

### Colony Formation Assay

To calculate the respective surviving fractions (SF) after γ radiation (0, 1, 2, 4, 6 and 8 G), a standard colony-forming assay was performed, as previously described (Rave-Fränk et al. 2007). Briefly, cells were exposed to γ-radiation and incubated for 7 days. Next, cells were fixed with 70 % ethanol and stained with Mayer’s hemalum (Cat#1.09249.0500, Merck). Non-irradiated cultures were used for normalization. Colonies with >50 cells were scored as survivors. The experiments were performed three independent times, each time in triplicate, and calculated as the median. To validate statistical differences between control and treatment groups, analysis of variance (ANOVA: Two-Factor with Replication) was performed using Microsoft Excel software (version 2016 MSO). P-values < 0.05 were scored as significant.

### Accumulation and recovery of γH2AX signals

Cells were seeded on 96-well, black/clear, tissue culture treated plates (Corning Falcon cat# 353219). To allow cell attachment, cells were incubated at 37°C 5% CO_2_ for 18 hours. Next, cells were treated with 250 ng/ml NCS for 30 minutes and allowed to recover in fresh medium for the time points indicated. Cells were then fixed in 4% formaldehyde in PBS for 20 minutes and permeabilized with 0.2% Triton-X-100 in PBS for 20 minutes. Cells were washed with PBS and blocked with 10% FBS in PBS for 30 minutes. The plates were incubated with primary antibodies for 2 hours. Following three PBS wash steps; cells were incubated for 60 minutes with the respective secondary antibodies and 1:5000 DAPI. The γH2AX signal intensities were measured on a Celigo 4 Channel Imaging Cytometer (Nexlecom Bioscience). The image was quantified using the Celigo Software. Images were taken using a Nikon Eclipse Ti2-E Inverted microscope, Plan Apoλ oil immersion objective 100x NA1.45 WD=0.13 (Nikon Instruments Inc) and processed using NIS-Elements AR 5.02.00 (Nikon Instruments Inc), and FIJI (ImageJ) for the preparation of figures. These experiments were performed at least 3 independent times, each time with duplicates and calculated as the mean.

### Statistical analysis

All graphs were generated using the PRISM8 software, statistical analysis was carried out by unpaired or paired two-tailed t-test, one-way or two-way ANOVA. For protein analysis from immunoblots, ImageStudio software was used after developing of the membranes by LI-COR. Bands were quantified and related to the respective loading control. Mean was calculated by the PRISM software from at least three independent experiments and represented together with SEM error bars. For Colony Formation and Cell Viability assays, measurements from three independent experiments with technical triplicates per data point are represented as mean ± SEM. For microscopy images, dots from PLAs or dots from foci formation assays were quantified with DUO-LINK software.

## Acknowledgments

The authors thank Ulrike Möller and Bettina Mayer for excellent technical assistance. RGF is thankful to Frauke Melchior for critical reading of this manuscript. This work was supported by the Deutsche Forschungsgemeinschaft Grant 2234/1-3

## REFERENCES

Ajami, K. et al., 2004. Dipeptidyl peptidase 9 has two forms, a broad tissue distribution, cytoplasmic localization and DPIV-like peptidase activity. Biochimica et Biophysica Acta (BBA) - Gene Structure and Expression, 1679(1), pp.18–28.

Bachmair, A., Finley, D. & Varshavsky, A., 1986. In vivo half-life of a protein is a function of its amino-terminal residue. Science, 234(4773), pp.179–186.

Carreira, A. & Kowalczykowski, S.C., 2011. Two classes of BRC repeats in BRCA2 promote RAD51 nucleoprotein filament function by distinct mechanisms. Proceedings of the National Academy of Sciences of the United States of America, 108(26), pp.10448–10453.

Chen, C.-C. et al., 2018. Homology-Directed Repair and the Role of BRCA1, BRCA2, and Related Proteins in Genome Integrity and Cancer. Annual review of cancer biology, 2(1), pp.313–336.

Chen, S.-J. et al., 2017. An N-end rule pathway that recognizes proline and destroys gluconeogenic enzymes. Science, 355(6323), p.eaal3655.

Davies, A.A. et al., 2001. Role of BRCA2 in control of the RAD51 recombination and DNA repair protein. Molecular Cell, 7(2), pp.273–282.

Davies, O.R. & Pellegrini, L., 2007. Interaction with the BRCA2 C terminus protects RAD51-DNA filaments from disassembly by BRC repeats. Nature Structural & Molecular Biology, 14(6), pp.475–483.

de Vasconcelos, N.M. et al., 2019. DPP8/DPP9 inhibition elicits canonical Nlrp1b inflammasome hallmarks in murine macrophages. Life Science Alliance, 2(1), p.e201900313.

Dong, C. et al., 2018. Molecular basis of GID4-mediated recognition of degrons for the Pro/N-end rule pathway. Nature Chemical Biology, 14(5), pp.466–473.

Elia, A.E.H. et al., 2015. RFWD3-Dependent Ubiquitination of RPA Regulates Repair at Stalled Replication Forks. Molecular Cell, 60(2), pp.280–293.

Esashi, F. et al., 2007. Stabilization of RAD51 nucleoprotein filaments by the C-terminal region of BRCA2. Nature Structural & Molecular Biology, 14(6), pp.468–474.

Feeney, L. et al., 2017. RPA-Mediated Recruitment of the E3 Ligase RFWD3 Is Vital for Interstrand Crosslink Repair and Human Health. Molecular Cell, 66(5), pp.610–621.e4.

Galanty, Y. et al., 2012. RNF4, a SUMO-targeted ubiquitin E3 ligase, promotes DNA double-strand break repair. Genes & Development, 26(11), pp.1179–1195.

Gall, M.G. et al., 2013. Targeted inactivation of dipeptidyl peptidase 9 enzymatic activity causes mouse neonate lethality. G. Sotiropoulou, ed. PLoS ONE, 8(11), p.e78378.

Geiss-Friedlander, R. et al., 2009. The cytoplasmic peptidase DPP9 is rate-limiting for degradation of proline-containing peptides. Journal of Biological Chemistry, 284(40), pp.27211–27219.

Gong, Z. & Chen, J., 2011. E3 ligase RFWD3 participates in replication checkpoint control. Journal of Biological Chemistry, 286(25), pp.22308–22313.

Inano, S. et al., 2017. RFWD3-Mediated Ubiquitination Promotes Timely Removal of Both RPA and RAD51 from DNA Damage Sites to Facilitate Homologous Recombination. Molecular Cell, 66(5), pp.622–634.e8.

Jasin, M. & Rothstein, R., 2013. Repair of Strand Breaks by Homologous Recombination. Cold Spring Harbor Perspectives in Biology, 5(11), pp.a012740–a012740.

Jensen, R.B., Carreira, A. & Kowalczykowski, S.C., 2010. Purified human BRCA2 stimulates RAD51-mediated recombination. Nature, 467(7316), pp.678–683.

Johnson, D.C. et al., 2018. DPP8/DPP9 inhibitor-induced pyroptosis for treatment of acute myeloid leukemia. Nature Medicine, pp.1–12.

Justa-Schuch, D. et al., 2016. DPP9 is a novel component of the N-end rule pathway targeting the tyrosine kinase Syk. eLife, 5, p.14741.

Justa-Schuch, D., Möller, U. & Geiss-Friedlander, R., 2014. The amino terminus extension in the long dipeptidyl peptidase 9 isoform contains a nuclear localization signal targeting the active peptidase to the nucleus. Cellular and Molecular Life Sciences (CMLS), 71(18), pp.3611–3626.

Kari, V. et al., 2016. Loss of CHD1causes DNA repair defects and enhances prostate cancer therapeutic responsiveness. EMBO reports, 17(11), pp.1609–1623.

Krawczyk, P.M. et al., 2011. Mild hyperthermia inhibits homologous recombination, induces BRCA2 degradation, and sensitizes cancer cells to poly (ADP-ribose) polymerase-1 inhibition. Proceedings of the National Academy of Sciences of the United States of America, 108(24), pp.9851–9856.

Li, J. et al., 2006. DSS1 is required for the stability of BRCA2. Oncogene, 25(8), pp.1186–1194.

Liu, Jie et al., 2010. Human BRCA2 protein promotes RAD51 filament formation on RPA-covered single-stranded DNA. Nature Structural & Molecular Biology, 17(10), pp.1260–1262.

Liu, Jinping et al., 2017. Ubiquitin-specific protease 21 stabilizes BRCA2 to control DNA repair and tumor growth. Nature Communications, 8(1), pp.137–12.

Liu, Shangfeng et al., 2011. RING finger and WD repeat domain 3 (RFWD3) associates with replication protein A (RPA) and facilitates RPA-mediated DNA damage response. Journal of Biological Chemistry, 286(25), pp.22314–22322.

Lord, C.J. & Ashworth, A., 2016. BRCAness revisited. Nature Reviews Cancer, 16(2), pp.110–120.

Luo, K. et al., 2012. Sumoylation of MDC1 is important for proper DNA damage response. The EMBO Journal, 31(13), pp.3008–3019.

Magwood, A.C., Mundia, M.M. & Baker, M.D., 2012. High Levels of Wild-Type BRCA2 Suppress Homologous Recombination. Journal of Molecular Biology, 421(1), pp.38–53.

Mavaddat, N. et al., 2013. Cancer risks for BRCA1 and BRCA2 mutation carriers: results from prospective analysis of EMBRACE. JNCI Journal of the National Cancer Institute, 105(11), pp.812–822.

Melnykov, A., Chen, S.-J. & Varshavsky, A., 2019. Gid10 as an alternative N-recognin of the Pro/N-degron pathway. Proceedings of the National Academy of Sciences of the United States of America, 116(32), pp.15914–15923.

Menear, K.A. et al., 2008. 4-[3-(4-cyclopropanecarbonylpiperazine-1-carbonyl)-4-fluorobenzyl]-2H-phthalazin-1-one: a novel bioavailable inhibitor of poly(ADP-ribose) polymerase-1. Journal of Medicinal Chemistry, 51(20), pp.6581–6591.

Moynahan, M.E., Pierce, A.J. & Jasin, M., 2001. BRCA2 is required for homology-directed repair of chromosomal breaks. Molecular Cell, 7(2), pp.263–272.

Okondo, M.C. et al., 2016. dpp8 and dpp9 inhibition induces pro-caspase-1-dependent monocyte and macrophage pyroptosis. Nature Chemical Biology, pp.1–10.

Okondo, M.C. et al., 2018. Inhibition of Dpp8/9 Activates the Nlrp1b Inflammasome. Cell chemical biology.

Oliver, A.W. et al., 2009. Structural basis for recruitment of BRCA2 by PALB2. 10(9), pp.990–996.

Pellegrini, L. et al., 2002. Insights into DNA recombination from the structure of a RAD51– BRCA2 complex. Nature, 420(6913), pp.287–293.

Pilla, E. et al., 2012. A novel SUMO1-specific interacting motif in dipeptidyl peptidase 9 (DPP9) that is important for enzymatic regulation. Journal of Biological Chemistry, 287(53), pp.44320–44329.

Pilla, E. et al., 2013. The SUMO1-E67 interacting loop peptide is an allosteric inhibitor of the dipeptidyl peptidases 8 and 9. Journal of Biological Chemistry, 288(45), pp.32787–32796.

Rave-Fränk, M. et al., 2007. Comparison of the combined action of oxaliplatin or cisplatin and radiation in cervical and lung cancer cells. International journal of radiation biology, 83(1), pp.41–47.

Ross, B. et al., 2018. Structures and mechanism of dipeptidyl peptidases 8 and 9, important players in cellular homeostasis and cancer. Proceedings of the National Academy of Sciences of the United States of America, 115(7), pp.E1437–E1445.

Rottenberg, S. et al., 2008. High sensitivity of BRCA1-deficient mammary tumors to the PARP inhibitor AZD2281 alone and in combination with platinum drugs. Proceedings of the National Academy of Sciences, 105(44), pp.17079–17084.

Sánchez, H. et al., 2017. Architectural plasticity of human BRCA2-RAD51 complexes in DNA break repair. Nucleic Acids Research, 45(8), pp.4507–4518.

Schoenfeld, A.R. et al., 2004. BRCA2 is ubiquitinated in vivo and interacts with USP11, a deubiquitinating enzyme that exhibits prosurvival function in the cellular response to DNA damage. Molecular and Cellular Biology, 24(17), pp.7444–7455.

Scully, R. et al., 2019. DNA double-strand break repair-pathway choice in somatic mammalian cells. Nature Reviews Molecular Cell Biology, 40(7732), p.179.

Shahid, T. et al., 2014. Structure and mechanism of action of the BRCA2 breast cancer tumor suppressor. Nature Structural & Molecular Biology, 21(11), pp.962–968.

Smebye, M.L. et al., 2017. Involvement of DPP9 in gene fusions in serous ovarian carcinoma. pp.1–10.

Spagnuolo, P.A. et al., 2013. Inhibition of intracellular dipeptidyl peptidases 8 and 9 enhances parthenolide’s anti-leukemic activity. Leukemia, 27(6), pp.1236–1244.

Tang, Z. et al., 2017. Contribution of upregulated dipeptidyl peptidase 9 (DPP9) in promoting tumoregenicity, metastasis and the prediction of poor prognosis in non-small cell lung cancer (NSCLC). International Journal of Cancer, 140(7), pp.1620–1632.

Thorslund, T. et al., 2010. The breast cancer tumor suppressor BRCA2 promotes the specific targeting of RAD51 to single-stranded DNA. Nature Publishing Group, 17(10), pp.1263–1265.

Torella, J.P. et al., 2014. Unique nucleotide sequence-guided assembly of repetitive DNA parts for synthetic biology applications. Nature Protocols, 9(9), pp.2075–2089.

Turinetto, V. & Giachino, C., 2015. Multiple facets of histone variant H2AX: a DNA double-strand-break marker with several biological functions. Nucleic Acids Research, 43(5), pp.2489–2498.

Varshavsky, A., 2019a. N-degron and C-degron pathways of protein degradation. Proceedings of the National Academy of Sciences, 116(2), pp.358–366.

Varshavsky, A., 2005. Ubiquitin fusion technique and related methods. Methods in enzymology, 399, pp.777–799.

Velkova, A. et al., 2014. Identification of Filamin A as a BRCA1-interacting protein required for efficient DNA repair. Cell Cycle, 9(7), pp.1421–1433.

Wang, S.-C. et al., 2002. Inhibition of cancer cell growth by BRCA2. Cancer Research, 62(5), pp.1311–1314.

Waumans, Y. et al., 2015. The Dipeptidyl Peptidase Family, Prolyl Oligopeptidase, and Prolyl Carboxypeptidase in the Immune System and Inflammatory Disease, Including Atherosclerosis. Frontiers in Immunology, 6(1), pp.1786–18.

Wilson, C.H. et al., 2013. Identifying Natural Substrates for Dipeptidyl Peptidases 8 and 9 Using Terminal Amine Isotopic Labeling of Substrates (TAILS) Reveals in VivoRoles in Cellular Homeostasis and Energy Metabolism. Journal of Biological Chemistry, 288(20), pp.13936–13949.

Wu, J.-J. et al., 2009. Biochemistry, pharmacokinetics, and toxicology of a potent and selective DPP8/9 inhibitor. Biochemical Pharmacology, pp.1–8.

Xia, B. et al., 2006. Control of BRCA2 Cellular and Clinical Functions by a Nuclear Partner, PALB2. Molecular Cell, 22(6), pp.719–729.

Yang, H. et al., 2005. The BRCA2 homologue Brh2 nucleates RAD51 filament formation at a dsDNA–ssDNA junction. Nature, 433(7026), pp.653–657.

Yin, Y. et al., 2012. SUMO-targeted ubiquitin E3 ligase RNF4 is required for the response of human cells to DNA damage. Genes & Development, 26(11), pp.1196–1208.

Yuan, Y. & Shen, Z., 2001a. Interaction with BRCA2 suggests a role for filamin-1 (hsFLNa) in DNA damage response. Journal of Biological Chemistry, 276(51), pp.48318–48324.

Yuan, Y. & Shen, Z.Y., 2001b. Interaction with BRCA2 suggests a role for filamin-1 (hsFLNa) in DNA damage response. The Journal of biological chemistry, 276(51), pp.48318–48324.

Yue, J. et al., 2012. Filamin-A as a marker and target for DNA damage based cancer therapy. DNA Repair, 11(2), pp.192–200.

Yue, J. et al., 2009. The Cytoskeleton Protein Filamin-A Is Required for an Efficient Recombinational DNA Double Strand Break Repair. Cancer Research, 69(20), pp.7978–7985.

Yue, J., Huhn, S. & Shen, Z., 2013. Complex roles of filamin-A mediated cytoskeleton network in cancer progression. Cell & Bioscience, 3(1), pp.7–12.

Zhang, F. et al., 2009. PALB2 Functionally Connects the Breast Cancer Susceptibility Proteins BRCA1 and BRCA2. Molecular Cancer Research, 7(7), pp.1110–1118.

Zhang, H., Chen, Y., et al., 2015. Dipeptidyl peptidase 9 subcellular localization and a role in cell adhesion involving focal adhesion kinase and paxillin. BBA -Molecular Cell Research, 1853(2), pp.470–480.

Zhang, H., Maqsudi, S., et al., 2015. Identification of novel dipeptidyl peptidase 9 substrates by two-dimensional differential in-gel electrophoresis. The FEBS journal, 282(19), pp.3737–3757.

